# Distinct networks of expressed genes are associated with neophobia in the hippocampus of male and female Eurasian tree sparrows (*Passer montanus*)

**DOI:** 10.1101/2025.09.04.674257

**Authors:** S.E. Lipshutz, A. B. Bentz, E. B. Cochran, K. J. Krajcir, M. G. Kimball, C. R. Lattin

## Abstract

Neophobia, avoidance of novel stimuli, is an ecologically and evolutionarily relevant behavioral trait that varies among individuals and across species. Especially among wild animals, the neuromolecular mechanisms underlying individual variation in neophobia have not been well characterized. We examined three neophobic behaviors in captive female and male Eurasian tree sparrows (*Passer montanus*) from a wild population introduced to the USA in 1870: responses towards novel objects, novel foods, and repeated presentations of the same initially novel object. We compared transcriptomic patterns associated with neophobia in three brain regions, the striatum, dorsal hippocampus, and rostral hippocampus, using differential expression and co-expression network analyses. We found that the striatum and hippocampus had distinct transcriptomic profiles, as did the rostral and caudal subregions of the hippocampus, supporting recent hypotheses that these subregions are functionally specialized. Despite the absence of sex differences in neophobic behaviors, neophobia-associated gene modules revealed sex-specific patterns within brain regions. For females, neophobic behaviors more strongly correlated with gene modules in the caudal hippocampus, a region involved in stress and anxiety, whereas for males, neophobic behaviors correlated with gene modules in the rostral hippocampus, a region that may play a larger role in spatial cognition. These modules exhibited significant overlap, suggesting that neophobic behaviors in both females and males are driven by shared neurobiological mechanisms, though they exhibit sex-specific patterns of brain region localization. Further, this work highlights the importance of examining both male and female animals in neurobiological research.

## Introduction

Fearful behavior towards novel stimuli, or neophobia, is found in many animal species, and can be observed in response to new foods, objects, and environments (Crane & Ferrari, 2017; Kimball & Lattin, 2023b). While an aversive reaction to a dangerous new stimulus can help ensure an animal’s survival, a fearful response to novelty may prevent access to new foods, nesting sites, or other beneficial resources (Greenberg & Mettke-Hofmann, 2001). Neophobia has become increasingly relevant for many wild animals as their environments undergo anthropogenic changes. Research suggests that urban animals are less neophobic than their rural counterparts (Breck et al., 2019; Huang et al., 2020; Jarjour et al., 2019; Tryjanowski et al., 2016) and even within urban sites, individuals that experienced more anthropogenic disturbances were less neophobic than those encountering fewer disruptions (Grunst et al., 2019). Neophobia has also been tied to invasion success, where reduced neophobia may help invasive species exploit novel resources (Candler & Bernal, 2015; Cohen et al., 2020; Martin & Fitzgerald, 2005). However, despite its ecological relevance and the wide variation in neophobia observed across individuals, populations, and species, the molecular mechanisms underlying this behavior are not well understood. Therefore, there is a clear need for more studies investigating neurobiological differences between neophobic and non-neophobic individuals.

Much of what is currently known about the neurobiological basis of neophobia comes from laboratory rodent models. Although lab-bred rats and mice typically show a preference for novel over familiar stimuli, they do exhibit neophobia towards some - but not all - novel tastes and when feeding in a novel environment (hyponeophagia) (Olszewski et al., 2014; Roman et al., 2009; Shephard et al., 1982; Stehberg et al., 2011). Rodent research has identified several neurobiological mediators in the glutamate, serotonin, and norepinephrine systems involved in these types of feeding neophobia (Ramírez-Lugo et al., 2015; Royet et al., 1983; Simpson et al., 2011; Steketee et al., 1992). However, because these neophobic responses are context- and stimulus-specific, it is unclear how generalizable these results are to the more widespread neophobia typically observed in wild animals (Crane & Ferrari, 2017). Further, because lab animals tend to be highly inbred, they do not show the wide within-species variation in neophobia observed in many wild populations. Research in wild-caught or outbred animals is necessary to reveal the neuroendocrine basis for individual variation in neophobia.

Transcriptomic approaches enable the unbiased identification of gene expression patterns associated with behavior, extending the scope beyond the handful of neurotransmitters and hormones typically investigated with more targeted approaches (Bell & Robinson, 2011; Bengston et al., 2018; Calisi & MacManes, 2015). Transcriptomics has been particularly useful in revealing what brain regions are involved in behaviors with wide inter-individual variation, like neophobia. For example, a recent transcriptome study in the invasive house sparrow (*Passer domesticus*) (Lattin et al., 2022) found that neophobic and non-neophobic individuals had distinct patterns of constitutive differential gene expression in brain regions that included the hippocampus (a region linked to spatial cognition, novelty responses, and memory) and the striatum (which is involved in reward and learning) (Güntürkün, 2005; Medina & Reiner, 1995; Rose & Colombo, 2005; Shinohara & Yasoshima, 2021). Differences between neophobic and non-neophobic sparrows were especially pronounced in the hippocampus, suggesting this region may be particularly important in neophobia (Lattin et al., 2022), a conclusion further supported by a study showing high neuronal activity in the hippocampus after exposure to novel objects (Kimball et al., 2022). However, this transcriptome work was performed in only one sex, leaving uncertainty about sex-specific mechanisms.

Although many species - including house sparrows - show no sex differences in neophobia (Schaffer et al., 2021; Kelly et al., 2020; Krajcir et al., 2024), it is not necessarily the case that the same neurobiological mechanisms regulate the same behaviors in males and females. The sexes may converge upon similar behaviors despite divergent neurobiological processes, due to different life history pressures or endocrine phenotypes (McCarthy et al., 2012). Neurobiological differences between females and males may emerge in several different ways. For instance, many sex differences have been found in hippocampal anatomy in mammals (Brivio et al., 2020; Yagi & Galea, 2019). In laboratory mice, females have greater dendritic spine density and heightened gene expression responses to receptor knockouts in the hippocampus than males, likely driven by differences in steroid hormone levels (Oakley et al., 2024; Shors et al., 2001). In birds, female brown-headed cowbirds (*Molothrus ater*) have a larger hippocampus and greater neurogenesis than males, potentially associated with females’ brood parasite life history strategy (Guigueno et al., 2016; Sherry et al., 1993). However, food-caching black-capped chickadees (*Poecile atricapillus*) do not show sex differences in spatial memory or hippocampal size (Petersen & Sherry, 1996), and recent genome-wide association studies revealed little effect of sex on the genetic contribution to chickadee spatial cognition (Semenov et al., 2024). Thus, the inclusion of both female and male subjects in biological research has the potential to reveal sex-dependent biological processes (Klein et al., 2015; Shansky & Murphy, 2021).

In the present study, we examined potential links between neophobia and constitutive gene expression in the brain of Eurasian tree sparrows (*Passer montanus*), an invasive songbird introduced to the USA in 1870 (Widmann, 1889). Birds are among the taxa with the highest levels of neophobia (Crane & Ferrari, 2017), and like their congener the house sparrow, Eurasian tree sparrows have wide and repeatable inter-individual variation in neophobia (Krajcir et al., 2024). Because different types of neophobia tests can sometimes show varying results (Dusang et al., 2025; Kimball & Lattin, 2023a), neophobia was assessed using three different behavioral assays: sparrows’ responses towards novel objects, novel foods, and repeated exposures of the same initially novel object. Three weeks after neophobia behavior tests were complete, we compared gene expression patterns in three different brain regions (striatum, rostral hippocampus, and caudal hippocampus) in males and females in relation to their behavioral performance using differential expression and co-expression network analyses. We divided the hippocampus into subregions because recent evidence suggests that the avian hippocampus shows specialization along a rostral-caudal axis, similar to that of the dorsal and ventral hippocampus in rodents (Fanselow & Dong, 2010; Gualtieri et al., 2019; Smulders, 2017). We predicted that neophobic and non-neophobic individuals would show differences in constitutive gene expression across the three brain regions, with distinct patterns reflecting their functional specializations. These patterns could emerge via differences in expression patterns, module composition or structure, or biological processes, suggesting that there may be multiple biological routes to the same behavioral outcome. Based on past work showing novel object exposure increased neuronal activity in the caudal but not rostral portions of the house sparrow hippocampus (Kimball et al., 2022), we expected the strongest effects of neophobia on expression in the caudal hippocampus. In the same set of Eurasian tree sparrows, we previously found no sex differences in responses to any neophobia assays (Krajcir et al., 2024). However, although neophobia behaviors did not differ by sex, we predicted that the underlying transcriptomic patterns might show sex specificity.

## Materials and methods

### Bird capture and behavioral trials

All animal work was approved by the Louisiana State University Institutional Animal Care and Use Committee (protocol #10-2021), and Eurasian tree sparrows were collected under an Illinois State Scientific Collecting Permit. Sparrow capture, transport, housing, and euthanasia followed bird research guidelines from the Ornithological Council (Fair et al., 2023). Adult Eurasian tree sparrows (n = 13 males, 11 females) were captured from Tazewell County, Illinois, United States in March 2023 using mist nets and nest box traps on private landowners’ property. Sparrows were transported to Louisiana State University overnight (to minimize stress) in a modified pet carrier with multiple food and water dishes and perches. Once birds arrived at Louisiana State University, they were placed in long-term housing cages in a vivarium facility on campus. The cages were supplied with multiple perches of differing materials, a dust bathing tray filled with sand, a plastic pine branch to use as a hide, water, and food (a mix of seeds, vitamin-supplemented food pellets, and grit). Birds were acclimated to captivity for three weeks before beginning behavior trials. Neophobia behavior trials are further detailed in Krajcir et al. (2024). Birds were doubly housed until three days before behavioral trials. Because Eurasian tree sparrows are not sexually dimorphic, we did not know sex at the time of paired housing. We paired individuals randomly, so some were in same-sex and some were in mixed-sex pairs. None of the pairs showed aggression during shared housing. During experimental trials, birds were singly housed to avoid any effects of social interactions on neophobia (Kelly et al., 2020).

Sparrows were cared for daily by the research team and animal husbandry staff, and weighed every two weeks to ensure body mass was maintained.

Neophobia in captive Eurasian tree sparrows was quantified using the same methods previously used in house sparrows (Kimball et al., 2022). Briefly, food was removed 30 min before lights out in the room, and trials conducted the following morning 30 min after lights on. Neophobia was quantified as latency to first approach and feed from a food dish presented either with a novel object (a purple plastic egg, white partial cover, red-painted dish, yellow pipe cleaners, or blinking light) or a novel food item (chopped kiwi, shredded cheese, peanut butter, or fruit-flavored cereal) during 1-h trials. Multiple types of novel objects and foods were used to ensure we measured individuals’ general response to novelty, rather than specific responses to one particular stimulus, which may be a type of pseudoreplication (Kimball and Lattin 2023b). Control approach and feeding times were determined by presenting birds with their normal food dish filled with familiar foods. Control trials were used to ensure that behavioral responses were specific to the novel stimulus, rather than a generalized aversion to the testing procedure; indeed, all sparrows ate relatively quickly when only familiar foods were present during control trials (on average 214 ± 224 s to approach and 254 ± 232 s to feed).

Food neophobia, object neophobia, and habituation to repeated exposures of an initially novel object were tested. Object neophobia was tested during week one (n = 3 novel object trials n = 2 control trials per bird, presented in random order), food neophobia was tested during week two (n = 4 novel food trials and n = 2 control trials per bird, presented in random order), and object habituation was tested during week three (n = 4 trials using the same initially novel object and n = 1 control trial per bird, with the control trial first). There is some debate in the literature about what measure best reflects an animal’s neophobia: its responses to novel stimuli alone or the difference in its responses to novel compared to control stimuli (e.g., Many Birds Project 2025). Because we found that average responses to novel objects and foods and the difference from average control responses were correlated (R***^2^*** > 0.71, p < 0.0001 for all measures), we used sparrows’ average responses to novel stimuli for ease of interpretation. All six behavioral measures of neophobia (average latency to approach the food dish when novel objects were present, average latency to feed when novel objects were present, average latency to approach the food dish when it contained novel food, average latency to eat novel food, average latency to approach the food dish during novel object habituation trials, and average latency to feed during novel object habituation trials) were highly and significantly correlated among individuals when measured as a group, as well as when males and females were examined separately, except for novel food approach and novel food feeding in females (Table S1). To correlate different neophobia-related behaviors with gene expression (see WGCNA below), we averaged among trials for novel objects, food, and object habituation for each individual.

### RNAseq tissue preparation and sample collection

Three weeks after neophobia trials ended, sparrows were euthanized using an overdose of isoflurane anesthesia followed by rapid decapitation. Our goal was to assess differences in constitutive gene expression in the brains of neophobic and non-neophobic birds, rather than differences due to behavioral trials. This three week period was chosen because this is the typical length of time we use to habituate wild animals to new conditions such as lab housing, and because there is good evidence that this length of time is sufficient from some physiological changes to adjust (Dickens and Romero 2009). After decapitation, brains were removed, flash-frozen on powdered dry ice, and stored at -80°C until sectioned coronally on a cryostat (Cryostar NX50, Thermo Fisher). Tissue was embedded in Optimal Cutting Temperature compound (VWR 95057-838) in block molds and placed in the cryostat to be sliced at -22°C. Slices were mounted directly onto slides (Superfrost Plus, VWR 48311-703) in two alternating series of 50 μm and 200 μm slices (Lattin et al., 2022). The 50 μm slides were dried overnight at 4°C, stained with thionin, which stains Nissl substance and DNA, and coverslipped. The 200 μm slides were chilled on dry ice and stored at -80°C until tissue punching.

Brain tissue punches were taken from the three target regions: striatum, caudal hippocampus, and rostral hippocampus (Figure 1A). Target brain slices for punching in the 200 μm slides were determined using landmarks in the adjacent 50 μm thionin-stained slides, using published songbird brain atlases as guides (Nixdorf-Bergweiler & Bischof, 2007; Stokes et al., 1974). Striatum punches came from slices that also contained the quinto-frontal tract, which corresponds to plates 3 and 4 from the Stokes canary atlas (1974). Two 1 mm diameter punches (Fine Science Tools 18035–01, 15 G) were taken from each hemisphere of the striatum on one target slice (i.e., four total punches), ∼3 mm from the rostral tip of the brain. Rostral hippocampus was punched on slices where the septomesencephalic tract (TrSM) was evident, ∼5 mm into the brain, which corresponds to plate 9 from the Stokes canary atlas (1974). Ten 0.5 mm diameter punches (Fine Science Tools 18035-50, 19G) were collected from both hemispheres of the rostral hippocampus across three consecutive brain slices. Caudal hippocampus was punched from slices where the cerebellum first came in, ∼8 mm into the brain, which corresponds to plates 20 and 21 from the Stokes canary atlas (1974). Tissue was collected from both hemispheres of four consecutive slices using the 0.5 mm diameter tissue punch (10-16 punches total). All punches from the same target brain region and subject were combined in sterile D/RNAse Free 1.6 mL centrifuge tubes and stored at -80°C until RNA extraction. Tissue punching tools were sterilized between subjects and brain regions using D/RNAse Free solution (Cole-Parmer EW-78901-43) followed by submersion in 95% molecular-grade ethanol, DEPC-treated water, and 95% molecular-grade ethanol.

**Figure 1.**
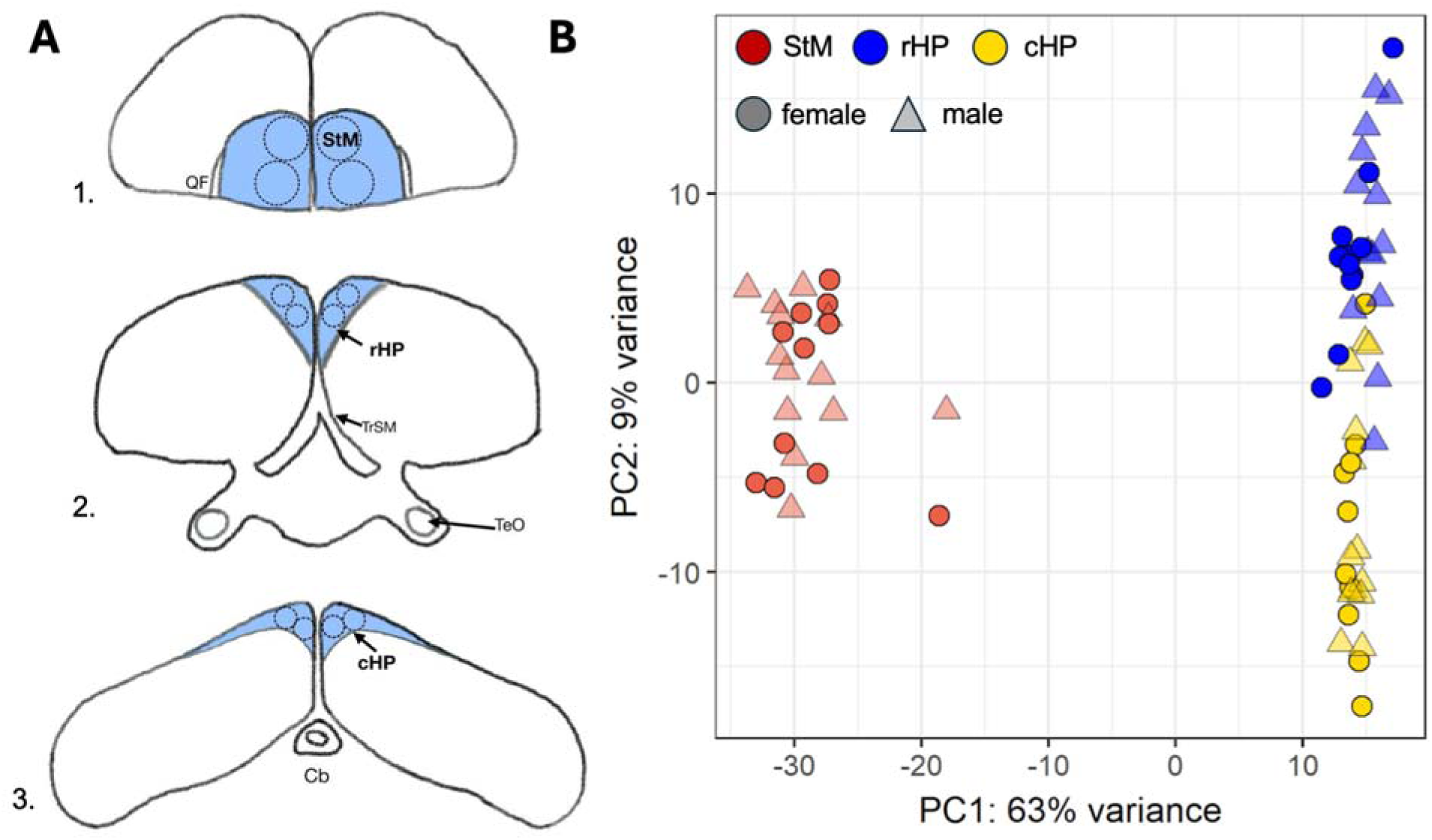
(A) Anatomical landmarks in hand-drawn coronal sections of Eurasian tree sparrow brains depicting the three target regions (shown in blue) as well as the relative size and location of the punches (dashed circles). The top image (1) is the most anterior slice and depicts the striatum (StM) and its landmark, the quinto-frontal tract (QF). The middle image (2) depicts the rostral hippocampus (rHP) along with its target landmark, the septomesencephalic tract (TrSM), as well as a rough depiction of the optic tectum (TeO). The bottom image (3) is the most posterior of the drawings, and depicts the caudal hippocampus (cHP), along with its target landmark, the cerebellum (Cb) at its first emergence. (B) Principal component analysis of gene expression patterns, showing the first two principal components (PC), which explain 72% of the variance. Points represent male (triangles) and female (circles) sparrows across three brain regions: striatum (red), caudal hippocampus (yellow), and rostral hippocampus (blue). Sample size = 13 males and 11 females per region.

### RNA extraction and library preparation

RNeasy Lipid Tissue Mini Kits (Qiagen; 1023539) were used to extract RNA from brain tissue and RNA concentrations and quality assessed using a Bioanalyzer system (Agilent 5400). Some samples initially resulted in RNA concentrations too low for library preparation.

Therefore, 13 samples were evaporated down in a sterile incubator until target concentrations for library preparation were achieved. The incubator was sterilized with D/RNase free and 100% molecular-grade ethanol before being brought to 45°C. Samples were evaporated for no more than 3 h. Final sample quality was assessed using the Bioanalyzer, and average total RIN score for all samples was 8.6 (range: 6.7-9.7).

### RNA sequencing, alignment, and mapping

We submitted total RNA to Novogene for sequencing using 150 bp paired-end sequencing (NovaSeq X Plus system) to generate an average of ∼52.4 million reads per sample (Table S2). After filtering to remove adapters and low-quality nucleotides, average reads per sample was ∼49.7 million. The error rate of sequencing for all samples was 0.01 and the G-C percentage was 48%.We used the zebra finch genome (GCA_003957565.4) as a reference (Rhie et al., 2021), as it is the most closely related species with a well-annotated genome (Hauber, Louder, & Griffith, 2021; Kumar et al., 2022). We aligned reads using HISAT2 (Mortazavi et al., 2008), for a 28.6% (range: 25.7-30.7%) mapping rate, with roughly ∼14 million reads per sample. We used StringTie to assemble transcripts (Pertea et al., 2015). We ran a principal component analysis (PCA) to visualize transcriptomic patterns across brain regions and sexes (Figure 1B).

### Identification of differentially expressed genes

We first identified sex- and region-specific genes that were differentially expressed between individuals showing the most consistent behavioral traits. We categorized individuals as “consistently neophobic” or “consistently non-neophobic” if their latency to approach or feed was above (more neophobic) or below (less neophobic) the median for novel object, novel object habituation, and novel food trials. Across all 6 behavioral trials, there were 7 individuals (n = 4 males; n = 3 females) that were consistently neophobic and 7 individuals (n = 4 males; n = 3 females) that were consistently non-neophobic (Figure S4). We acknowledge this binary approach reduced sample size, but it allowed us to focus on the most pronounced differences in gene expression associated with neophobia. Although latencies across trials were highly correlated, behavioral responses may have varied in non-binary ways; therefore, we also used gene co-expression networks to analyze the full spectrum of individual variation in each trial (see below).

Differentially expressed genes (DEGs) and normalized values were determined using DESeq2 v.1.42.0 in R (Love et al., 2014). Transcripts with < 10 counts in the smallest group sample size (n = 3 for females and n = 4 for males) were filtered out (n = 8,888 transcripts retained for female caudal hippocampus; n = 8,976 male caudal hippocampus; n = 8,957 female rostral hippocampus; n = 9,180 male rostral hippocampus; n = 8,869 female striatum; n = 8,772 male striatum). We performed Wald tests for each gene comparing non-neophobic vs neophobic (reference) individuals by sex within each brain region. Log2 fold change estimates were shrunk using the *apeglm* function in DESeq2. P-values were corrected using Benjamini-Hochberg corrections and FDR ≤ 0.05 were considered differentially expressed.

DEGs were assessed for enrichment of biological process Gene Ontology (GO) terms using PANTHER (Mi et al. Thomas, 2019) with a Fisher’s Exact test and a cut-off of FDR ≤ 0.05. GO terms were further summarized with REVIGO (http://revigo.irb.hr/), which clusters GO terms based on semantic similarity (similarity threshold = 0.7).

### Construction of weighted gene co-expression networks

Next, we performed a weighted gene co-expression network analysis (WGCNA) in each brain region (n = 24 individuals; 11 females and 13 males) to determine how whole networks of putatively co-regulated genes, rather than individual genes, relate to the full range of variation in latency to approach and feed during trials (Langfelder & Horvath, 2008). We created networks within brain regions because gene expression patterns were more region- than sex-specific (Figure 1B). Using the normalized counts from DESeq2, we filtered out genes with low counts (< 10 norm counts in the smallest group size; n = 11 for females) and low variability (25% least variable transcripts) within the hippocampus and striatum (n = 6,640 transcripts retained in cHp and rHp; n = 6,528 in striatum). Sample clustering was performed to detect sample-level outliers. We then generated a signed hybrid network by selecting a soft threshold power (β) = 3 for the caudal hippocampus, β = 9 for the rostral hippocampus, and β = 3 for the striatum in accordance with scale-free topology (signed R^2^ > 0.80; Figure S1). We used a biweight mid-correlation (bicor) function and modules were calculated using a minimum module size of 30. A threshold of 0.25 (merging modules with a correlation ≥ 0.75) was used to merge modules in Dynamic Tree Cut (Figure S2). Genes not assigned to a module were classified to the color gray. All other gene modules were assigned a random color.

### Identifying gene networks of interest

Expression levels for each module were summarized by the first principal component (module eigengene), which we used to test for an association between modules and the average time to approach or feed in each behavioral assay using the bicor function. We examined these relationships across and within sexes. We additionally included sex and body size (wing chord, tarsus length, and post-captivity mass) to account for these potentially confounding variables. We assessed each trait-associated module (defined as p < 0.05 and bicor > |0.60|) for enrichment of biological process GO terms using PANTHER (Mi et al., 2019). We only assessed genes with a significant module membership (MM; the correlation between the gene expression profile and module eigengene), defined as p < 0.05 and MM > 0.60.

We identified intramodular hub genes in trait-associated modules, as these genes are potential drivers of the patterns highlighted in these modules. We defined intramodular hub genes as those with a high trait-based gene significance (the correlation between a gene’s expression profile and a trait; GS > |0.60|) for at least one module-related trait of interest and high module membership (MM > 0.80), with a threshold of p < 0.05 for both (Horvath & Dong, 2008). We additionally looked for overlap between intramodular hub genes and DEGs as these genes have strong trait-related expression, making them core candidate hub genes. We identified whether these genes were found in the most enriched GO terms and visualized networks using Cytoscape (Shannon et al., 2003). Networks were filtered to only show edges supported by protein-protein interaction (PPI) data from the Search Tool for the Retrieval of Interacting Genes/Proteins (STRING) online database (https://string-db.org/) (Szklarczyk et al., 2019), with a combined score threshold of 0.4. We calculated the degree score for each gene, and those in the top 10% were classified as PPI hub genes.

To assess functional similarity across the two hippocampal subregions, we looked for consensus between the caudal and rostral hippocampus network modules by assessing similarity in module structure and gene membership. Modules were considered overlapping if they shared a significant number of genes, as determined by Fisher’s exact test. This approach allowed us to identify modules representing similar transcriptional profiles across brain regions. We additionally examined whether neophobia-related expression patterns emerged across brain regions by performing a permutational multivariate analysis of variance (PERMANOVA). This multivariate approach assesses group differences in the combined eigengene profile instead of testing each module separately. We compared sample-level module eigengenes from all behavioral trait-related modules across sex, neophobia phenotype, and their interaction in consistently neophobic and non-neophobic sparrows (n = 3 neophobic and 4 non-neophobic males; n = 3 neophobic and 3 non-neophobic females) (Figure S4). Module eigengenes summarize the expression pattern of a module and reflect how strongly a sample aligns with the overall module profile, allowing comparisons of co-expression patterns across networks within samples. We used the “adonis2” function in the R package *vegan* (Oksanen et al., 2019), with Euclidean distance, and visualized sample differences using PCA.

## Results

### Distinct transcriptomic profiles by brain region

One of our goals was to examine whether Eurasian tree sparrows showed distinct patterns of constitutive gene expression in the striatum as well as in different parts of the hippocampus.

Out of 8,935 genes, the majority (89.1%) were expressed in all three brain regions, though each region had ∼100-200 uniquely expressed genes (Figure S3). A PCA revealed that the three brain regions had distinct transcriptomic profiles (Figure 1B). PC1 explained 63% of the variance and partitioned the striatum from both hippocampal regions, while PC2 explained 9% of the variance and partitioned the hippocampal regions. Overall, these data suggest that the caudal and rostral hippocampus had distinct transcriptomic profiles, but were more similar to each other than to the striatum.

### Differentially expressed genes in neophobic and non-neophobic individuals by region and sex

Another goal was to compare sex- and region-specific differences in constitutive gene expression in consistently neophobic and non-neophobic individuals. As published previously (Krajcir et al., 2024), there were no sex differences in neophobia behaviors (p > 0.45) (Figure S4). Neither sex nor neophobia phenotype affected sparrows’ captive body mass (p > 0.07 at all time points). The greatest differences in constitutive gene expression were observed in the caudal hippocampus of females, followed by the rostral hippocampus of males (Figure S5). In females, there were n = 257 DEGs between non-neophobic and neophobic individuals in the caudal hippocampus (Figure S6A), n = 2 DEGs in the rostral hippocampus, and n = 4 DEGs in the striatum. In males, there were n = 2 DEGs between non-neophobic and neophobic individuals in the caudal hippocampus, n = 64 DEGs in the rostral hippocampus (Figure S6B), and n = 6 DEGs in the striatum. Only four DEGs were present in multiple contrasts, including neurotensin (NTS) in the male striatum and female caudal hippocampus, LOC100225836 in female striatum and caudal hippocampus, NBEAL2 in female striatum and caudal hippocampus, and NAPG in male rostral hippocampus and female caudal hippocampus (for all DEGs see Table S3).

Significantly enriched biological processes were only identified for DEGs in the female caudal hippocampus (Table S5). DEGs upregulated in neophobic females relative to non-neophobic females were most significantly enriched for “cytoplasmic translation” and “ribosome biogenesis”, while DEGs upregulated in non-neophobic females were most enriched for “neuron cellular homeostasis” and “learning or memory”, as well as other processes like “cellular responses to BDNF” and “Gamma-aminobutyric acid (GABA) metabolic processes”. While no GO biological processes were found, several of the DEGs in other regions are involved in neuroendocrine processes (e.g., SCG2 and NTS in the male striatum and RIT2 in the male caudal hippocampus), neuron signaling (e.g., GRM2 in the female striatum and GRIK2 in male rostral hippocampus), and stress responsiveness (e.g., OPRK1 in female rostral hippocampus).

### Trait-associated modules in the caudal hippocampus

We also used co-expression network analyses to examine which genes in the hippocampus and striatum were most associated with the full range of neophobic or non-neophobic behaviors. Within the caudal hippocampus, WGCNA constructed 9 modules (Table S5), several of which demonstrated trait-related patterns in females but not males (Figure 2A). The blue module was positively associated with novel food and novel object habituation approach times in females, while the turquoise module was negatively associated with these traits (Figures 2A, 2B). Genes in the blue module were most enriched for terms linked to translation (e.g., “cytoplasmic translation” and “ribosome assembly”) and energy production (e.g., “aerobic respiration” and “proton motive force-driven mitochondrial ATP synthesis”) (Figure 3A; Table S6), suggesting co-regulated genes involved in these processes are linked to slower approach times to novel food items and slower habituation to novel objects in females. Several of the intramodular hub genes in the blue module were differentially expressed between consistently neophobic and non-neophobic females (n = 38 genes), of which 12 genes were also PPI hub genes (Figure S7A), and the overlap between these intramodular hub genes and differentially expressed genes was significant (Fisher’s exact test, p < 0.01). These core candidate hub genes in the blue module were linked to some of the most enriched pathways (Figure 3A), suggesting they could be drivers of these processes.

**Figure 2.**
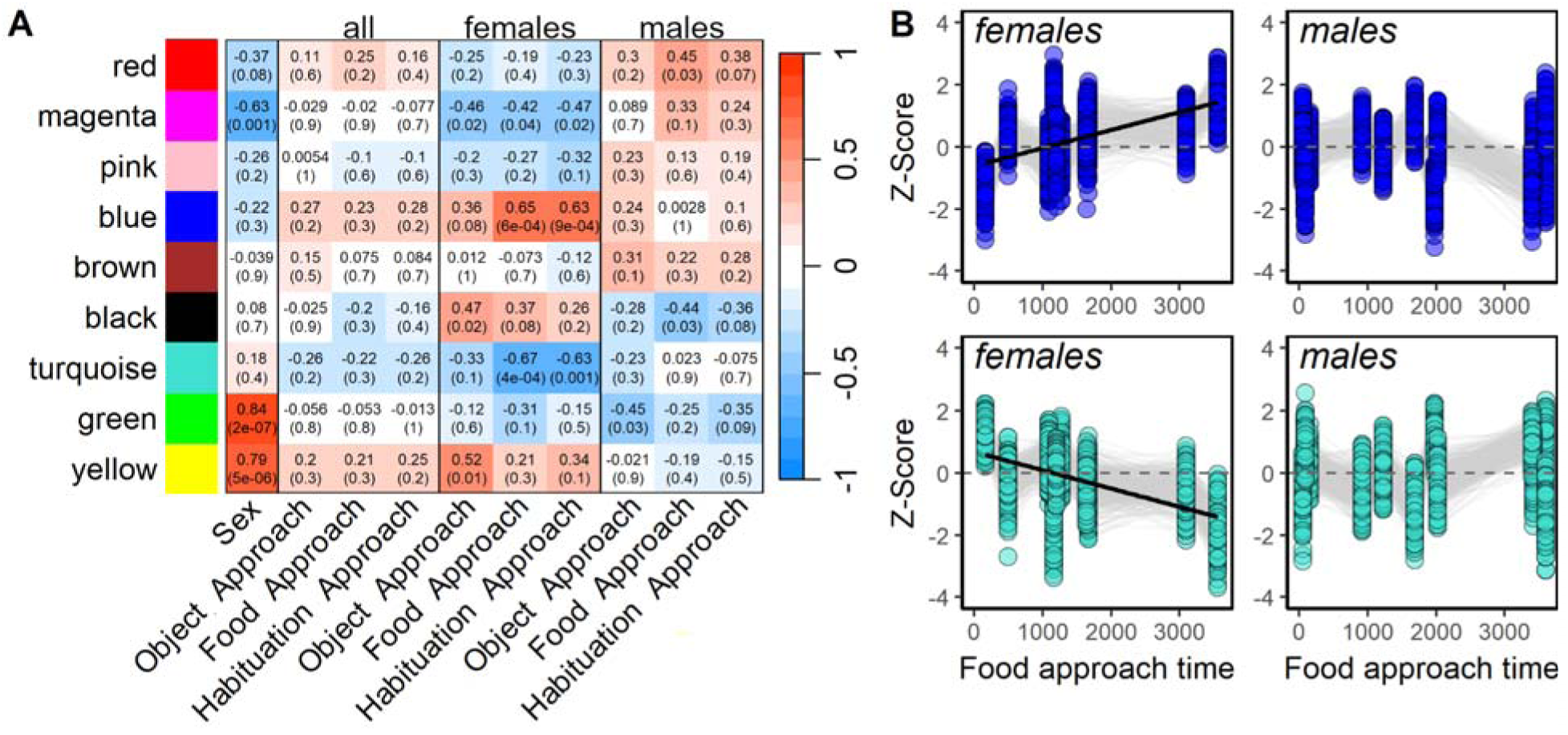
(A) Module-trait relationships in the Eurasian tree sparrow caudal hippocampus were identified using bicor tests in WGCNA. The colors of each gene module were assigned randomly. Associations are shown for sex (females as reference) and latency to approach in behavioral trials across and within sexes. “Object Approach” refers to the latency to approach a food dish with a novel object; “Food Approach” a food dish containing a novel food; and “Habituation Approach” a food dish with the same (initially novel) object over four days. For each individual, we averaged across multiple trials for each novel object, novel food, and habituation approach time measure. Correlation coefficients are shown with p-values in parentheses. The colors on the heatmap are determined by the magnitude and direction of the correlation coefficient, with darker shades of red indicating a stronger positive correlation and darker shades of blue indicating a stronger negative correlation. (B) Expression of genes in the blue (top) and turquoise (bottom) modules were significantly associated with neophobic behaviors in females but not males. Relative expression of genes with MM > 0.8 is shown across food approach times (in seconds). Gray lines connect genes across samples. Z-scores reflect standardized gene expression: Z = (x − μ)/σ, where μ and σ are the mean and standard deviation across all samples and x is the expression of the gene in a specific sample.

**Figure 3.**
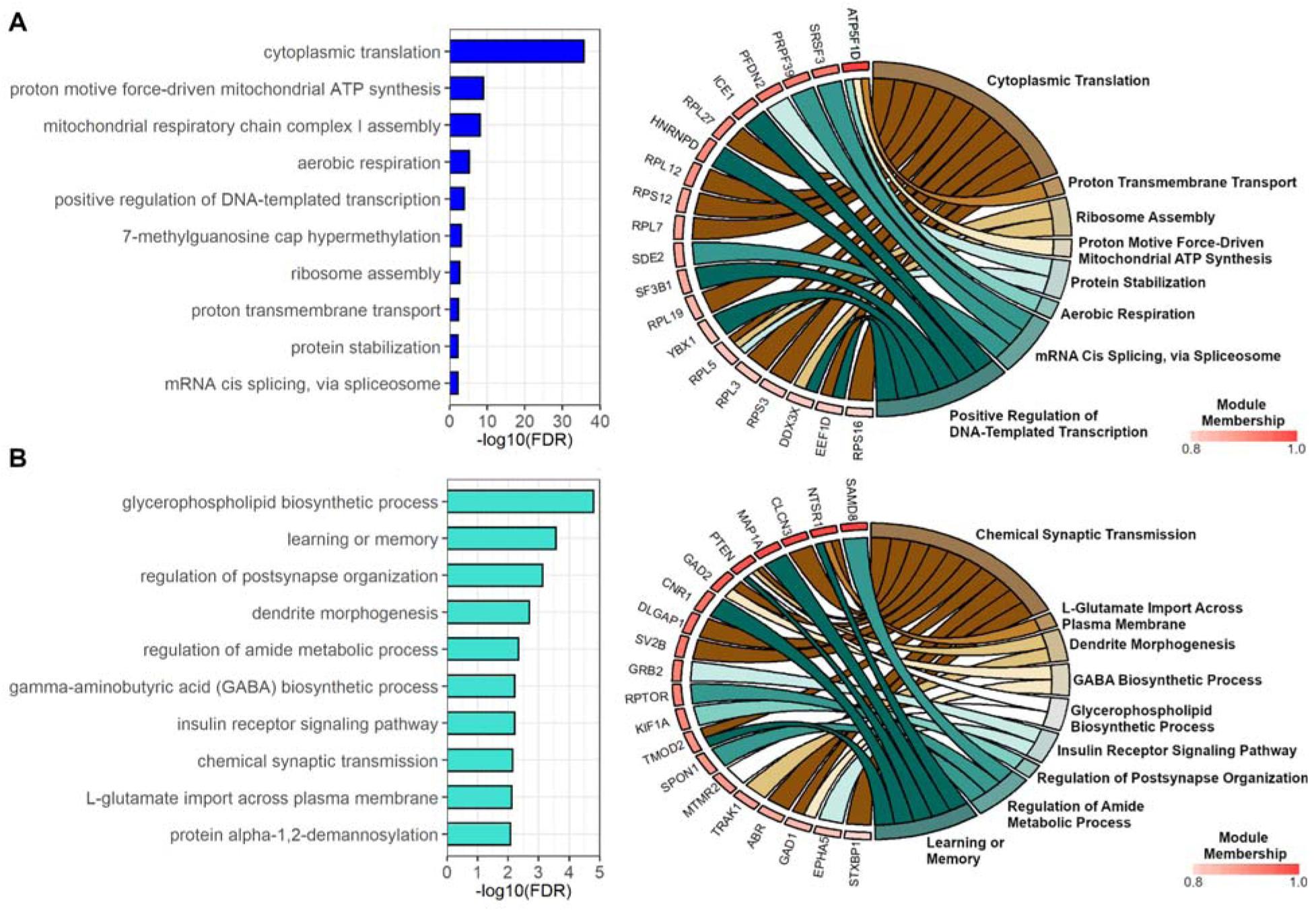
Barplot of the 10 most significant (i.e., lowest FDR) Gene Ontology (GO) biological processes and GOChord plot of associated genes in the (A) blue and (B) turquoise modules of the caudal hippocampus of Eurasian tree sparrows. Genes are linked to their assigned GO biological processes via colored ribbons and ordered according to their module membership. All genes shown are intramodular hubs and were differentially expressed between consistently neophobic and non-neophobic individuals.

Genes in the turquoise module were enriched for terms like “learning or memory”, “chemical synaptic transmission”, and “insulin receptor signaling pathway” (Figure 3B; Table S6), suggesting co-regulated genes involved in these processes are linked to faster approach times to novel food items and repeated presentations of the same initially novel object in females. Several of the intramodular hub genes in the turquoise module were also differentially expressed between consistently neophobic and non-neophobic females (n = 78 genes), of which 14 genes were also PPI hub genes (Figure S7B), and the overlap between these intramodular hub genes and differentially expressed genes was significant (Fisher’s exact test, p < 0.0001). These strong candidate hub genes were again linked to some of the most enriched biological processes (Figure 3B). Additional trait-related modules included the green and yellow modules that had genes expressed higher in males compared to females, and the magenta module that had genes with elevated expression in females relative to males (Figure S8). The magenta module was also negatively correlated with novel food feeding time (Figure S8). Functional enrichment results for all trait-associated modules can be found in Table S6.

### Trait-associated modules in the rostral hippocampus

In the rostral hippocampus, WGCNA constructed 8 modules (Table S7), several of which demonstrated trait-related patterns in males but not females (Figures 4A, S9). In males, the greenyellow module was positively associated with novel food approach time, while the black module was negatively associated with this behavior (Figure 4B). The purple module was negatively associated with novel object approach time and time to feed in the presence of a novel object in males (Figures 4A, S9). Genes in the greenyellow module were most enriched for terms related to cholesterol biosynthesis, steroid metabolism, and neurotransmitters (“GABA metabolic process” and “glutamate secretion”) (Figure 5A; Table S6), suggesting co-regulated genes involved in these processes are associated with slower approach times to novel food items in males. One intramodular hub gene in the greenyellow module was differentially expressed between consistently neophobic and non-neophobic males (SLC1A3), and it was also a PPI hub gene (Figure S10A). SLC1A3 is linked to the following enriched GO terms: “transepithelial transport”, “GABA metabolic process”, and “behavior”. Genes in the black module were enriched for terms related to transcription and post-transcriptional modifications, specifically mRNA splicing (Figure 5B; Table S6), suggesting co-regulated genes involved in these processes are linked to faster approach times to novel food items in males. Four intramodular hub genes in the black module were differentially expressed between consistently neophobic and non-neophobic males (MED13, SRSF7, RBBP6, and LOC100229022), of which SRSF7 was also a PPI hub gene (Figure S10B). SRSF7 is linked to the following enriched GO terms: “regulation of mRNA splicing, via spliceosome” and “mRNA cis splicing, via spliceosome”.

**Figure 4.**
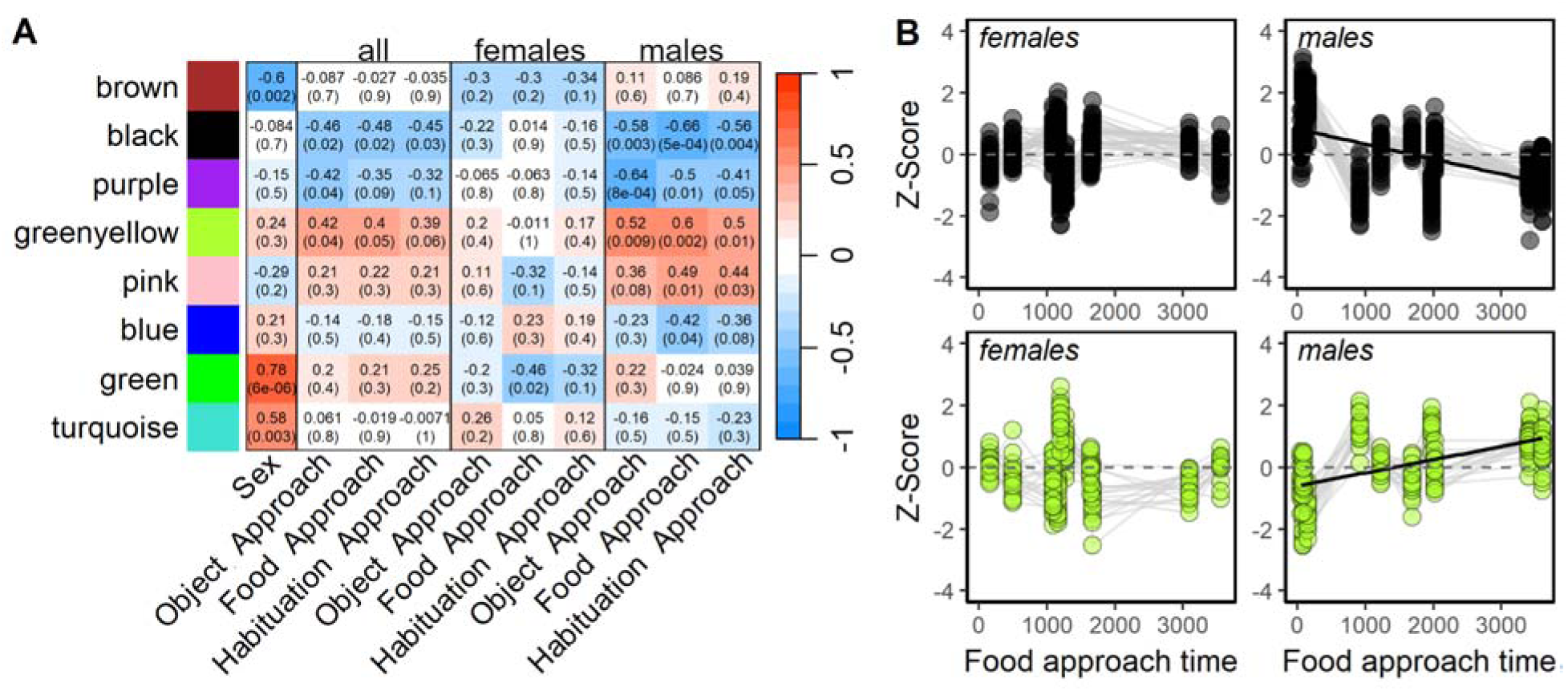
(A) Module-trait relationships in the sparrow rostral hippocampus were identified using bicor tests in WGCNA. The colors of each gene module were assigned randomly.

**Figure 5.**
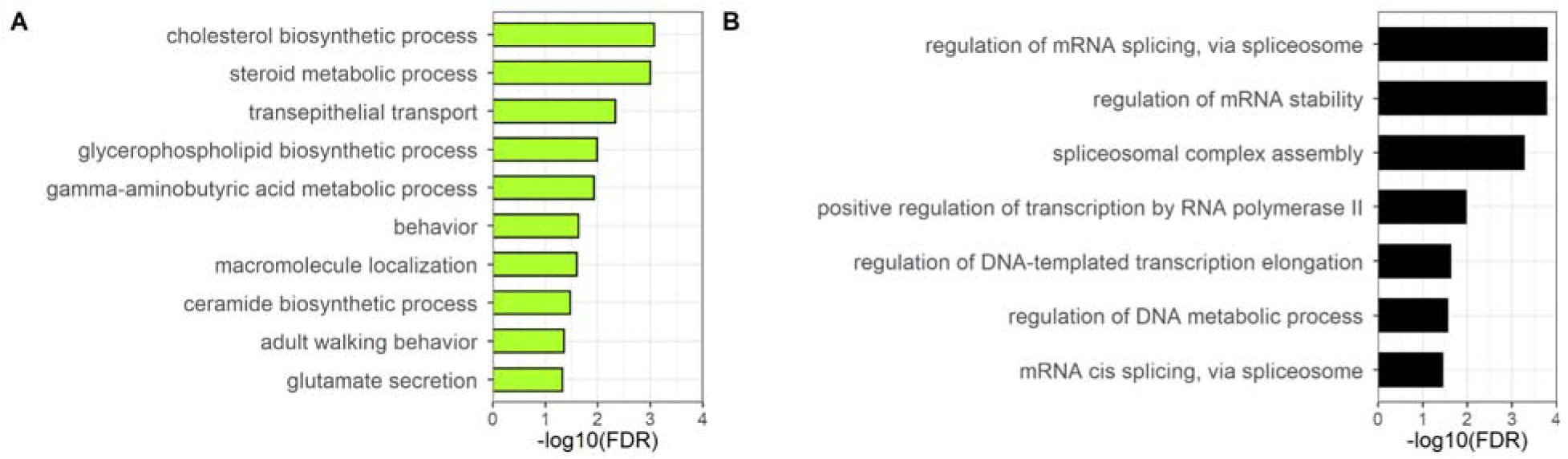
Barplot of the most significant (lowest FDR) Gene Ontology biological processes in the (A) greenyellow and (B) black modules of the rostral hippocampus of Eurasian tree sparrows.

Genes in the purple module were most enriched for a post-translational modification (i.e., “peptidyl-serine phosphorylation”; Table S6), suggesting this process could be linked to faster novel object approach and feeding times in males. One intramodular hub gene in the purple module was also differentially expressed between consistently neophobic and non-neophobic males (MACF1). Additional trait-related modules included the green and brown modules that had genes expressed higher in males and females, respectively (Figure S9). Of note, several critical stress-related genes, including CRH, NR3C1, and CRHR2 were assigned to the brown module. Functional enrichment results for all trait-associated modules can be found in Table S6.

Associations are shown for sex (females as reference) and latency to approach in behavioral trials across and within sexes. “Object Approach” refers to the latency to approach a food dish with a novel object; “Food Approach” a food dish containing a novel food; and “Habituation Approach” a food dish with the same (initially novel) object over four days. For each individual, we averaged across multiple trials for each novel object, novel food, and habituation approach time measure. Correlation coefficients are shown with p-values in parentheses. The colors on the heatmap are determined by the magnitude and direction of the correlation coefficient, with darker red shades indicating a stronger positive correlation and darker blue shades indicating a stronger negative correlation. (B) Expression of genes in the black (top) and greenyellow (bottom) modules were significantly associated with neophobic behaviors in males but not females. Relative expression of genes with MM > 0.8 is shown across food approach times (in seconds). Gray lines connect genes across samples. Z-scores reflect standardized gene expression: Z = (x − μ)/σ, where μ and σ are the mean and standard deviation across all samples and x is the expression of the gene in a specific sample.

### Trait-associated modules in the striatum

One outlier sample was removed from the striatum dataset (Figure S11). WGCNA constructed 12 modules in the striatum (Table S8), several of which demonstrated trait-related patterns in both males and females (Figures S12A, S13). The lightcyan module was negatively associated with multiple behavioral metrics in males, while the midnight blue, purple, and black modules were positively associated with neophobia metrics in males. The brown and red modules were positively associated with neophobia behavioral metrics in females. The most significant relationships were between the brown module and female novel object habituation approach time and the lightcyan module and male novel food approach time (Figure S12B).

Genes in the brown module were most enriched for terms linked to cellular transport and organization (i.e., “regulation of vesicle-mediated transport”, “positive regulation of organelle organization”, and “regulation of catabolic processes”; Table S6), suggesting co-regulated genes involved in these processes are linked to greater neophobia in females. There was no overlap between DEGs and intramodular hub genes in the brown module. Genes in the lightcyan module had no enriched terms, but one intramodular hub gene (HTT) was also differentially expressed between consistently neophobic and non-neophobic males, and it was also a PPI hub gene (Figure S14). Genes in the midnight blue module were enriched for terms related to sensory development (i.e., “visual system development” and “inner ear auditory receptor cell differentiation”) and signaling (i.e., notch and BMP signaling pathway terms), suggesting genes linked to these processes are elevated in more neophobic males, but there was no overlap between DEGs and intramodular hub genes. The black module was also most enriched for sensory system terms (i.e., “trigeminal nerve structural organization”, “branchiomotor neuron axon guidance”, “G protein-coupled opioid receptor signaling pathway”, “sensory perception of pain”, “adult behavior”, and “sensory system development”) and signaling processes (i.e., “neuropeptide signaling pathway” and “regulation of neuronal synaptic plasticity”), but there was no overlap between DEGs and intramodular hub genes. Genes in the purple module had no enriched terms, but one intramodular hub gene was also differentially expressed between consistently neophobic and non-neophobic males (SCG2). The red module was most enriched for mRNA splicing, suggesting genes linked to this process were elevated in more neophobic females, but there was no overlap between DEGs and intramodular hub genes. Additional trait-related modules included the yellow module that had genes expressed higher in males than females. Functional enrichment results for all trait-associated modules can be found in Table S6.

### Patterns across brain regions

Motivated by initial observations of transcriptional similarity, and to better understand shared patterns across the two subregions of the hippocampus and their sex-specific associations, we examined consensus between rostral and caudal hippocampal modules. We found significant gene overlap between neophobia-associated modules in the female caudal hippocampus and male rostral hippocampus, but relationships between neophobia and module gene expression were often reversed in the two sexes (Figure S14). Significant overlap in trait-associated modules included the blue module in the caudal hippocampus and the black module in the rostral hippocampus, the turquoise module in the caudal hippocampus and the greenyellow module in the rostral hippocampus, and the yellow module in the caudal hippocampus and green module in the rostral hippocampus (Figure 6). The blue-black module overlap involved 278 genes (constituting 77.9% of the black module genes) that were positively correlated with novel food approach times in the female caudal hippocampus (Figure 2B), but negatively correlated with this trait in the male rostral hippocampus (Figure 4B; Figure S15A). Genes found in both modules that had a MM > 0.80 and were significantly associated with behavioral traits were linked to transcription and post-transcriptional (e.g., mRNA splicing) processes (Figure S15

**Figure 6.**
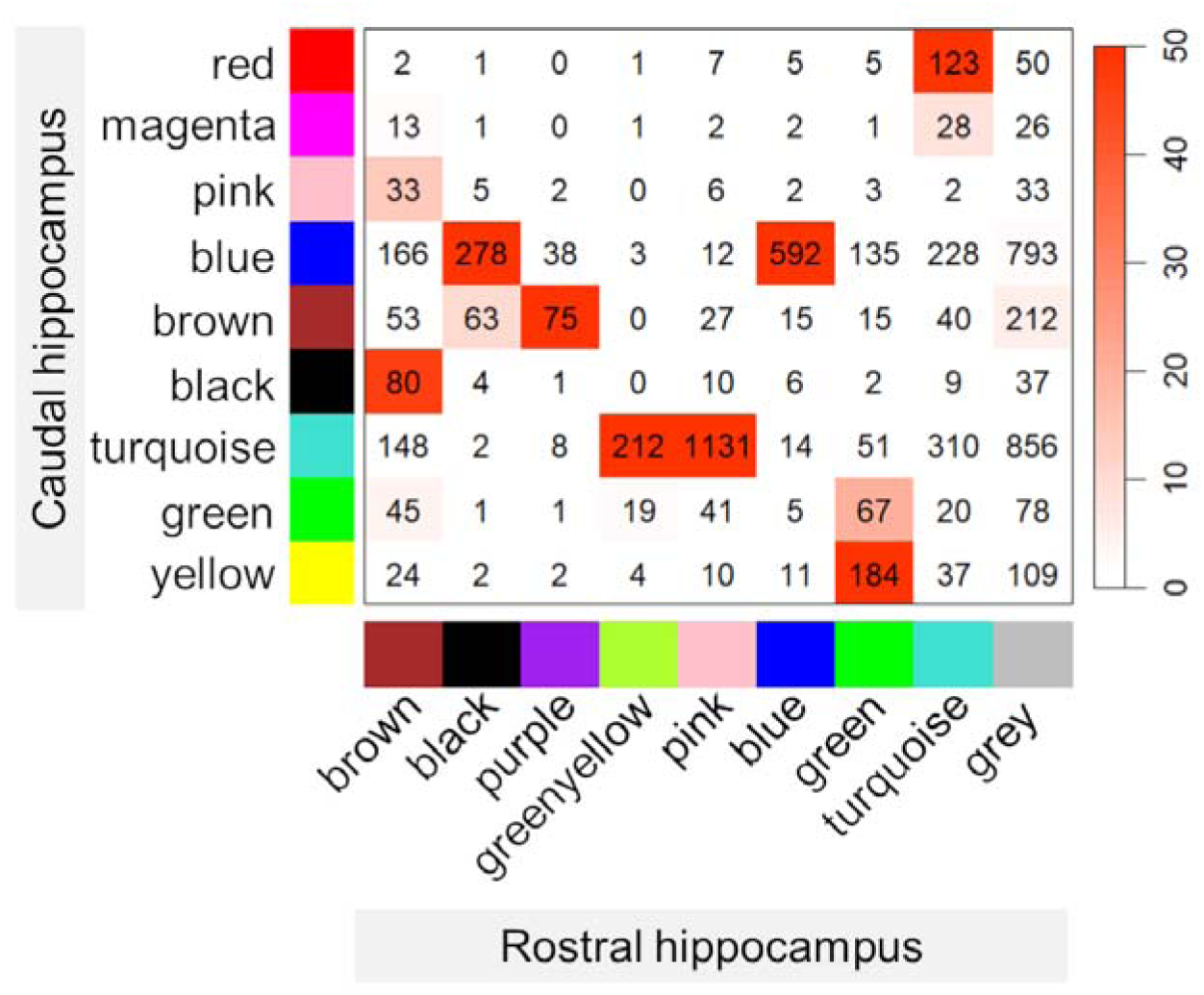
Correspondence between gene co-expression modules in the caudal and rostral hippocampus of Eurasian tree sparrows. Each row represents a module from the caudal hippocampus, and each column represents a module from the rostral hippocampus. Cell values indicate the number of transcripts shared between modules. Color intensity reflects the significance of module overlap; darker red color indicates a more significant overlap as calculated using Fisher’s exact test and shown as -log10(p-value).

A), suggesting that these processes may be elevated in the rostral hippocampus of less neophobic males and in the caudal hippocampus of more neophobic females. The turquoise-greenyellow overlap involved 212 genes (constituting 88.3% of the greenyellow module genes) that were negatively correlated with novel food approach times in the female caudal hippocampus (Figure 2B), but positively correlated with this trait in the male rostral hippocampus (Figure 4B; Figure S15A). Trait-related genes with high MM found in both modules were linked to chemical synaptic transmission and processes related to GABA and dopamine (Figure S16B), suggesting these processes may be elevated in the caudal hippocampus of less neophobic females and in the rostral hippocampus of more neophobic males. The yellow-green overlap involved 184 genes that were elevated in males in both brain regions.

We also examined whether neophobia-related expression patterns emerged across brain regions using the most consistently neophobic and non-neophobic individuals, as previously categorized for the differential expression analysis. We found that module eigengene profiles across brain regions significantly differed by behavioral phenotype. PERMANOVA results showed a significant main effect of neophobia (F_1,9_ = 3.44, p = 0.006), but not sex (F_1,9_ = 0.98, p = 0.45). There was a trend for the neophobia-by-sex interaction to be significant (neophobia by sex: F_1,9_ = 1.96, p = 0.08), thus we analyzed the principal component scores within each sex separately. Among males, more neophobic sparrows had significantly lower PC1 scores (F_1,5_ = 13.04, p = 0.015), but no difference in PC2 (F_1,5_ = 1.87, p = 0.23). Among females, more neophobic sparrows had significantly lower PC2 scores (F_1,4_ = 10.71, p = 0.03), but not PC1 (F_1,4_ = 0.003, p = 0.96). These differences were evident in the PCA, where the first two PCs explained 61.1% of the total variance in trait-related module eigengenes (Figure S17A). Lower PC1 scores in neophobic males reflected relatively lower expression of genes in the lightcyan module (striatum) and black module (rostral hippocampus), and higher expression in the greenyellow module (rostral hippocampus) (Figure S17B). In contrast, lower PC2 scores in neophobic females reflected higher expression in the brown module (striatum) and blue module (caudal hippocampus), and lower expression in the turquoise module (caudal hippocampus) (Figure S17 B). These results suggest that region-specific co-expression patterns are not isolated but instead co-occur within individuals in a behavior- and sex-dependent manner, likely reflecting coordinated transcriptional patterns across multiple brain regions.

## Discussion

We identified region-specific, constitutive transcriptomic patterns associated with neophobia in female and male Eurasian tree sparrows (*Passer montanus*). The striatum and hippocampus had distinct transcriptomic profiles, as did the rostral and caudal subregions of the hippocampus. The number of differentially expressed genes between neophobic and non-neophobic individuals varied across brain regions and sexes. However, the full complexity of the molecular mechanisms underlying neophobic behaviors was better captured using WGCNA, which revealed that neophobia was associated with similar gene modules between sexes, but their expression was localized to different hippocampal subregions in males and females. In females, neophobic behaviors were associated with gene modules in the caudal hippocampus tha were enriched for synaptic transmission, steroid metabolism, and post-transcriptional processes, whereas in males, neophobic behaviors were associated with similar gene modules in the rostral hippocampus. Altogether, these findings suggest that although neophobic behaviors are similar between sexes, they may be driven by region-specific neurobiological pathways. Below, we discuss these results and their implications for our understanding of the neurotranscriptomic mechanisms related to neophobia.

### Brain regions had distinct transcriptomic profiles, differentiating the caudal and rostral hippocampus

Brain region was a major driver of variation in gene expression. Though most genes were expressed in the striatum, caudal hippocampus, and rostral hippocampus, each region had ∼100-200 uniquely expressed genes. We found that the caudal and rostral hippocampus had distinct transcriptomic profiles, supporting the hypothesis that the avian hippocampus may show subregional specialization along a rostral-caudal axis, similar to the dorsal-ventral axis of the rodent hippocampus (Fanselow & Dong, 2010; Gualtieri et al., 2019; Smulders, 2017).

Specifically, the rostral hippocampus may play a larger role in spatial cognition, while the caudal hippocampus may be more involved in anxiety, stress, and avoidance responses (Madison et al., 2023). A recent study in black-capped chickadees also supports the functional specialization of these regions in songbirds, as the caudal hippocampus received more inputs from amygdalar regions involved in emotional responses, whereas the rostral hippocampus received more inputs from the thalamus, which plays a key role in relaying sensory and motor information (Applegate et al., 2023). To our knowledge, our study is the first to confirm that the rostral and caudal hippocampus express different transcriptomic profiles in birds. Future transcriptomic comparisons between the rostral and caudal subregions can further clarify whether different parts of the hippocampus are specialized for different functions in birds as well as mammals.

The striatum was most differentiated from the caudal and rostral hippocampus, as would be predicted based on its distinct structure and functions. Though subregional specializations of the striatum have not been well-explored in birds (Mezey & Csillag, 2002), the striatum of mammals is typically divided into the dorsal striatum, encompassing the caudate and putamen, and the ventral striatum, which includes the nucleus accumbens and the olfactory tubercle (Voorn et al., 2004). Future work could explore the transcriptomic profiles of striatal structures in birds as a means to better characterize potential striatal subregions.

### Neophobia-associated gene modules differed by brain region and sex

While prior work with this same group of Eurasian tree sparrows (as well as studies of the congeneric house sparrow) found no sex difference in neophobic behaviors (Ensminger et al., 2012; Kelly et al., 2020; Krajcir et al., 2024), we identified sex-specific patterns of neophobia-associated gene expression. Specifically, neophobic behaviors correlated with gene expression in the caudal hippocampus of females and the rostral hippocampus of males. Both categorical and continuous comparisons of gene expression related to neophobia revealed this sex-by-region interaction. Though more work is needed to understand the drivers of sex-specific gene expression in Eurasian tree sparrows, one hypothesis is that not only does the hippocampus show subregional specialization in birds, but that neural processes in these subregions vary by sex.

Notably, our finding in birds aligns with numerous studies from rodents showing that cognitive performance-related behaviors are associated with neurogenesis in the ventral hippocampus of females, and in the dorsal hippocampus of males (Dalla et al. 2009, Yagi et al. 2016, Yagi & Galea 2019). In rats, there are numerous sex differences in hippocampal anatomy that include dendritic spine density in the CA1 subregion (Shors et al., 2001; Woolley et al., 1990), morphology of CA3 pyramidal neurons (Juraska et al., 1989), and dendritic branching in the dentate gyrus (Juraska et al., 1985). Parallel differences may exist in the hippocampus of male and female birds, but this remains to be determined. This rodent work, together with the present study, suggest that sex differences in hippocampal processes may be conserved across vertebrates.

Our findings highlight that neophobia is regulated by sex-specific regionalization of gene expression, and suggest putative candidates for genes and processes that may play a conserved role in neophobia across species. In the caudal hippocampus, the most neophobic females had higher expression of genes involved in energy production and transcription/mRNA splicing/translation (blue module). The latter processes could increase transcriptomic diversity and functional complexity (Mauger & Scheiffele, 2017), yielding more diverse molecular routes to respond to novel stimuli. In contrast, non-neophobic females had higher expression of genes involved in various forms of signaling activity (turquoise module). This included GO terms related to learning or memory, synaptic transmission, and GABA biosynthesis. GABA is a key inhibitory neurotransmitter, and its synthesis depends on glutamic acid decarboxylases (GAD1 and GAD2), both of which were differentially expressed hub genes. Expression of these enzymes is highly correlated with GABA levels and inhibitory synaptic neurotransmission (Dicken et al., 2015; Lee et al., 2019), which shape neuronal activity and contribute to contextual learning in the hippocampus (Lovett-Barron et al., 2014). Additional differentially expressed hub genes included NTSR1 and CNR1, which are implicated in hippocampal learning and memory (Li et al., 2011; Ruiz-Contreras et al., 2013), as well as in modulating stress responses and anxiety-like behaviors (Kyriatzis et al., 2024; Martin et al., 2002). Thus, differences in the constitutive expression of these genes may influence how individuals respond to novel and potentially threatening stimuli. In the rostral hippocampus, the most neophobic males showed higher expression of genes involved in signaling activity, including GABA and steroids (greenyellow module), but lower expression of genes linked to transcription/mRNA splicing (black module).

Note that although we measured gene expression three weeks after behavioral trials ended, it is possible that exposure to novel stimuli initiated lasting effects on neuroplasticity that were still ongoing at the time of tissue collection. Alternatively, these genes may have differed in expression prior to the behavioral trials, and may explain some of the variation in neophobic responses. Regardless, these patterns suggest that neophobic males rely on specific signaling pathways rather than broad transcriptional flexibility in a hippocampal subregion involved in spatial processing, whereas non-neophobic females demonstrate a similar strategy but in a subregion associated with stress and anxiety.

Overall, our results suggest that basic neurobiological processes vary among individuals with different behavioral phenotypes, and that some parallel mechanisms may regulate neophobia in both sexes but in different brain regions and in opposite directions. This supports so-called ‘Type III’ sex differences (sensu (McCarthy et al., 2012)), where convergent behaviors are underlain by divergent neurobiological processes in females and males. What remains to be determined is how these individual and sex-specific differences in gene expression arise; is variation driven by innate neurobiological processes, or is it shaped by flexibly interacting with the environment? Activational and/or organizational effects of sex steroids could underlie these sex differences (Adkins-Regan, 2005). From a life-history standpoint, female songbirds are often more sensitive to stress-induced reproductive suppression, which could reflect sex-specific differences in brain regions involved in responding to potential stressors, like the hippocampus (Calisi et al., 2018; Hirschenhauser et al., 2003; Ouyang et al., 2012; Schmidt et al., 2014). As more studies (finally!) include both males and females, we encourage sex contextualist approaches (Pape et al., 2024; Richardson, 2022; Smiley et al., 2024) that assess the possibility of sex-related variation in neurobiology even in the absence of behavior differences, as we would not have discovered these sex-specific associations between neophobia and neural gene expression otherwise.

## Acknowledgements

We thank the landowners who provided property access to capture sparrows as well as field stations and their staff for housing for our personnel and sparrows during field work. Additionally, we thank members of the Lattin lab for their assistance with data collection, including Tosha Kelly, Keegan Stansberry, Ayushi Patel, Blake Dusang, Danna Masri, and Will Frazier. This work was funded by the US National Science Foundation CAREER grant IOS-2237423 (C.R.L.), CAREER grant IOS-2439462 (A.B.B.), IntBIO grant DBI-2316364 (S.E.L.), as well as support from Louisiana State University, Duke University, and the University of Oklahoma.

