## Supplemental File for "Distinct networks of expressed genes are associated with neophobia in the hippocampus of male and female Eurasian tree sparrows (*Passer montanus*)"

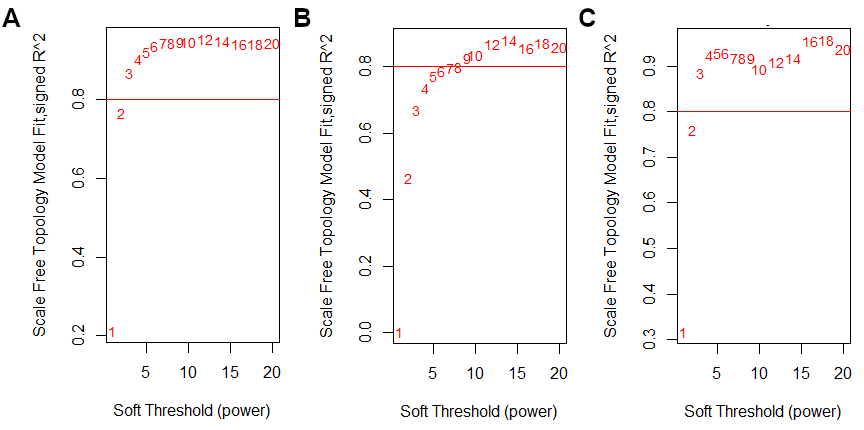

**Figure S1**. Scale-free topology for (A) caudal hippocampus, (B) rostral hippocampus, and (C) striatum.

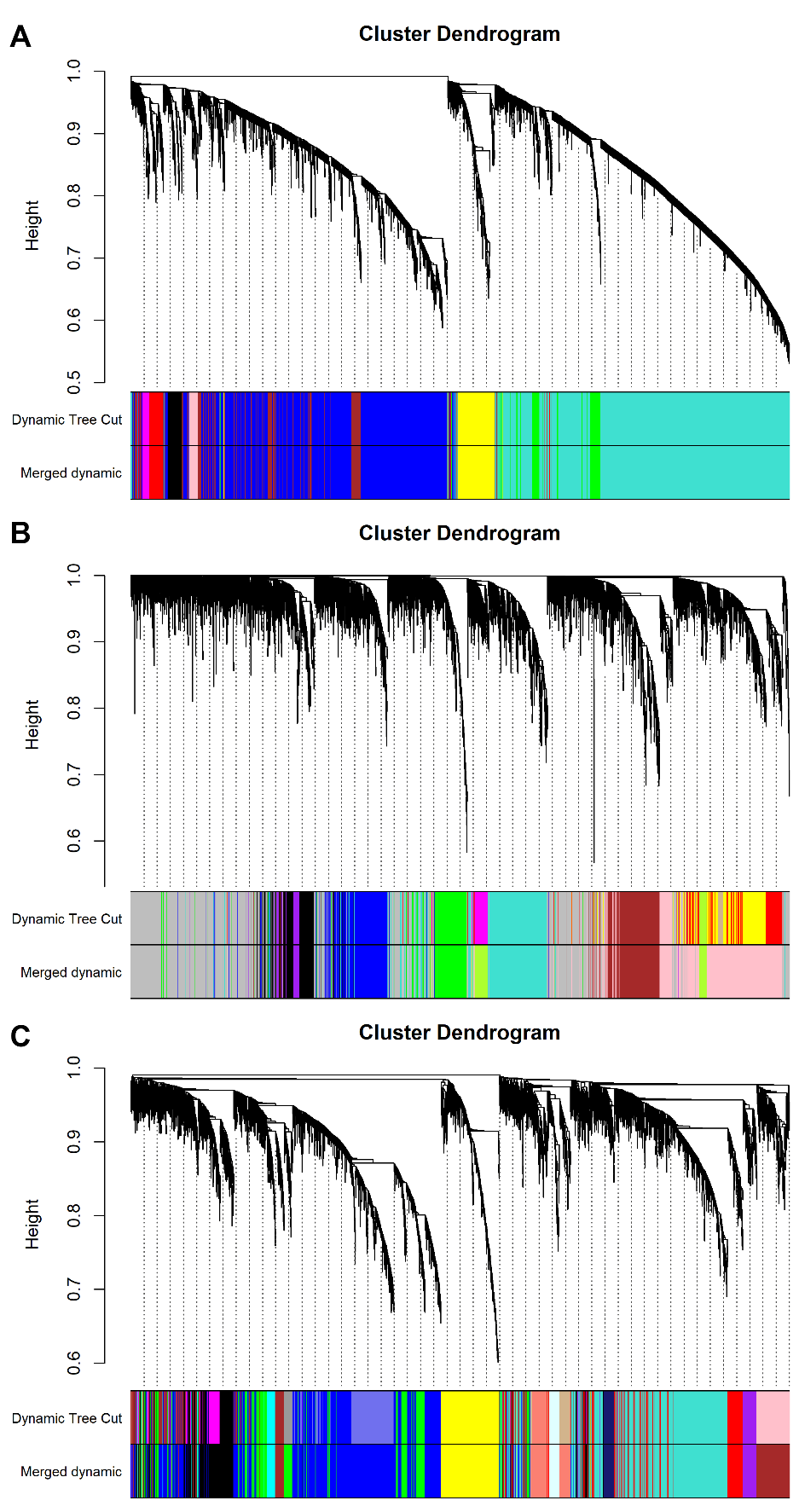

**Figure S2**. Gene dendrogram obtained by average linkage hierarchical clustering for (A) caudal hippocampus, (B) rostral hippocampus, and (C) striatum. The color row underneath the dendrogram shows the module assignment determined by Dynamic Tree Cut.

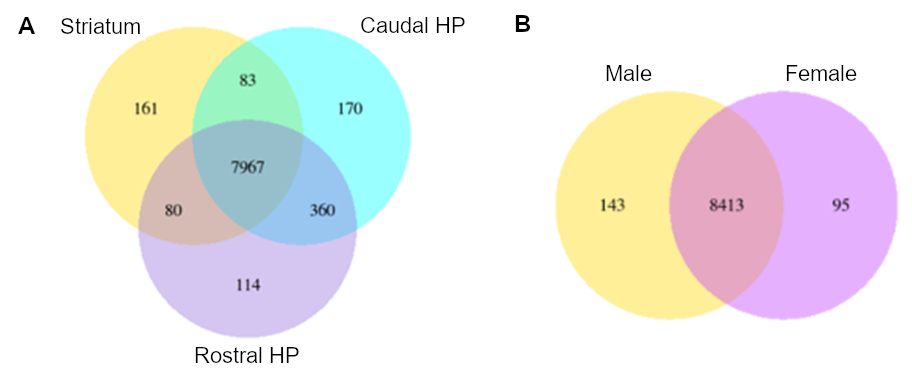

**Figure S3.** Venn diagrams of gene expression in Eurasian tree sparrows. (A) Overlap in expressed genes across three brain regions: striatum, rostral hippocampus, and caudal hippocampus (n = 24 per region). (B) Overlap in gene expression between the sexes. HP = hippocampus

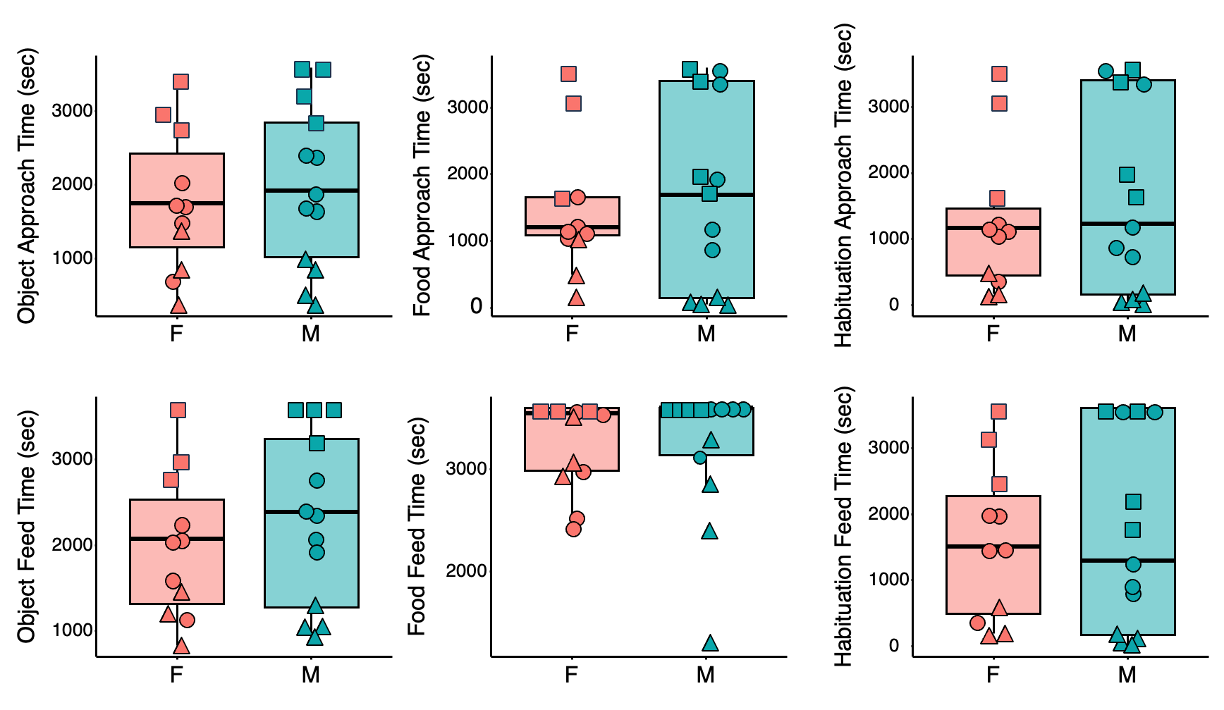

**Figure S4.** Behavioral responses to novel foods, objects, and habituation to the same initially novel object for approach latency (top row) and feed latency (bottom row) in seconds, in female (salmon) and male (turquoise) Eurasian tree sparrows. Consistently neophobic individuals are indicated by squares, whereas consistently non-neophobic individuals are indicated by triangles. Multiple trials were averaged for each individual. There were no sex differences in neophobia responses for any behavioral measure (p > 0.45) (Krajcir et al. 2024).

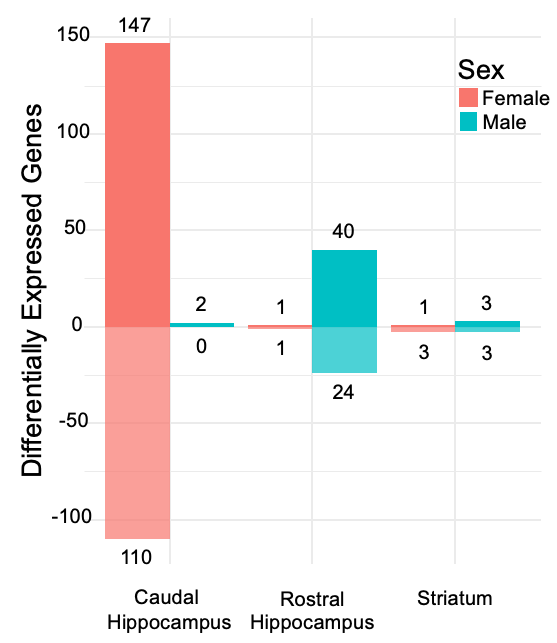

**Figure S5**. Barplots showing the number of differentially expressed genes between neophobic and non-neophobic sparrows across brain regions. The female contrast included n = 3 consistently neophobic and n = 3 consistently non-neophobic individuals. The male contrast included n = 4 consistently neophobic individuals and n = 4 consistently non-neophobic individuals. Bars are grouped by brain region and colored by sex: females (salmon), males (turquoise). For each group, bars extend upward for genes upregulated in non-neophobic individuals and downward for genes downregulated relative to neophobic individuals, based on log2 fold change direction.

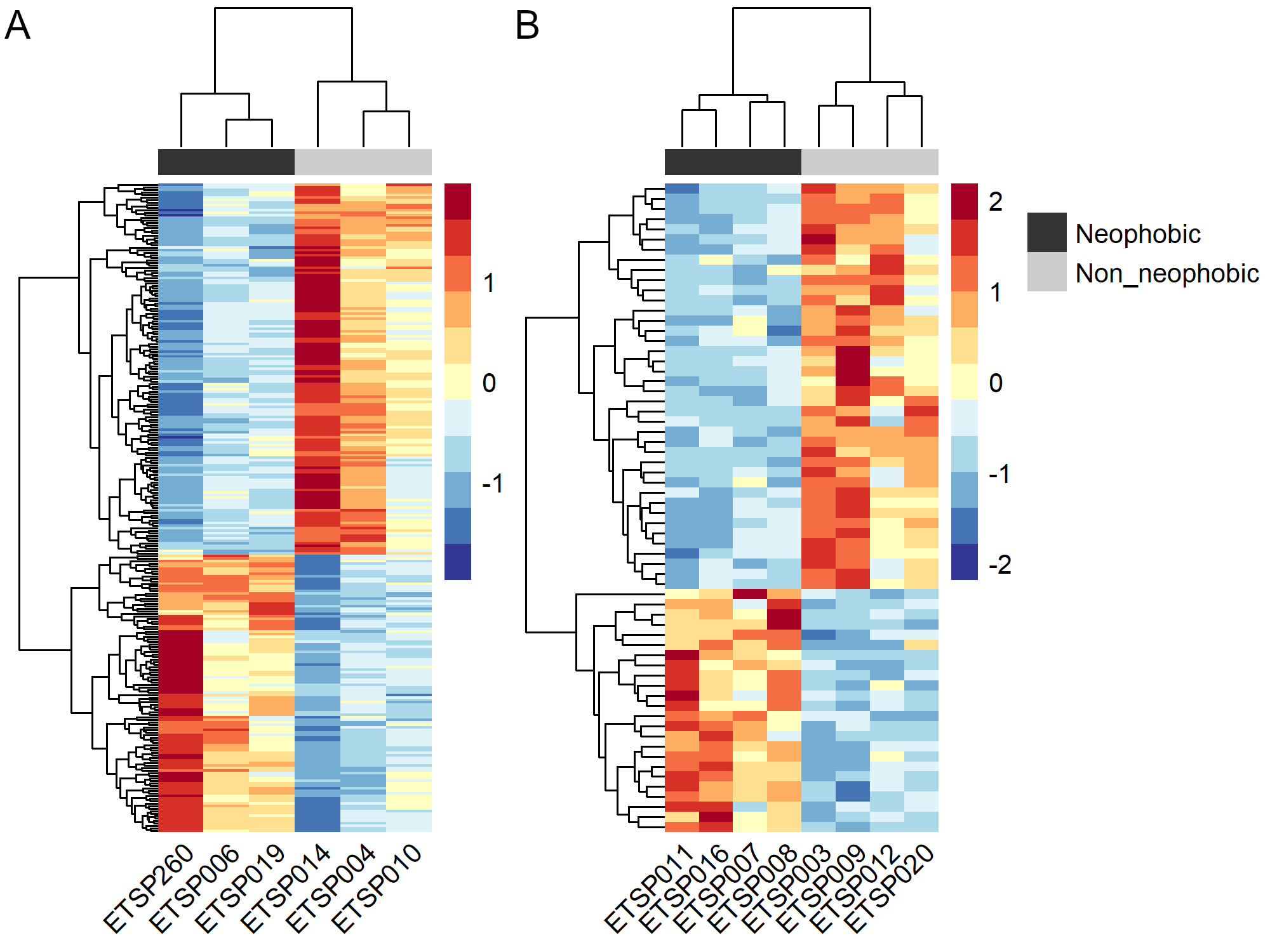

**Figure S6**. Heatmaps of differentially expressed genes between neophobic (black) and non-neophobic (gray) sparrows in the (A) female caudal hippocampus and (B) male rostral hippocampus. Colors indicate relative expression levels, with red denoting genes upregulated and blue denoting genes downregulated in non-neophobic individuals (relative to neophobic). Expression values are scaled across samples.

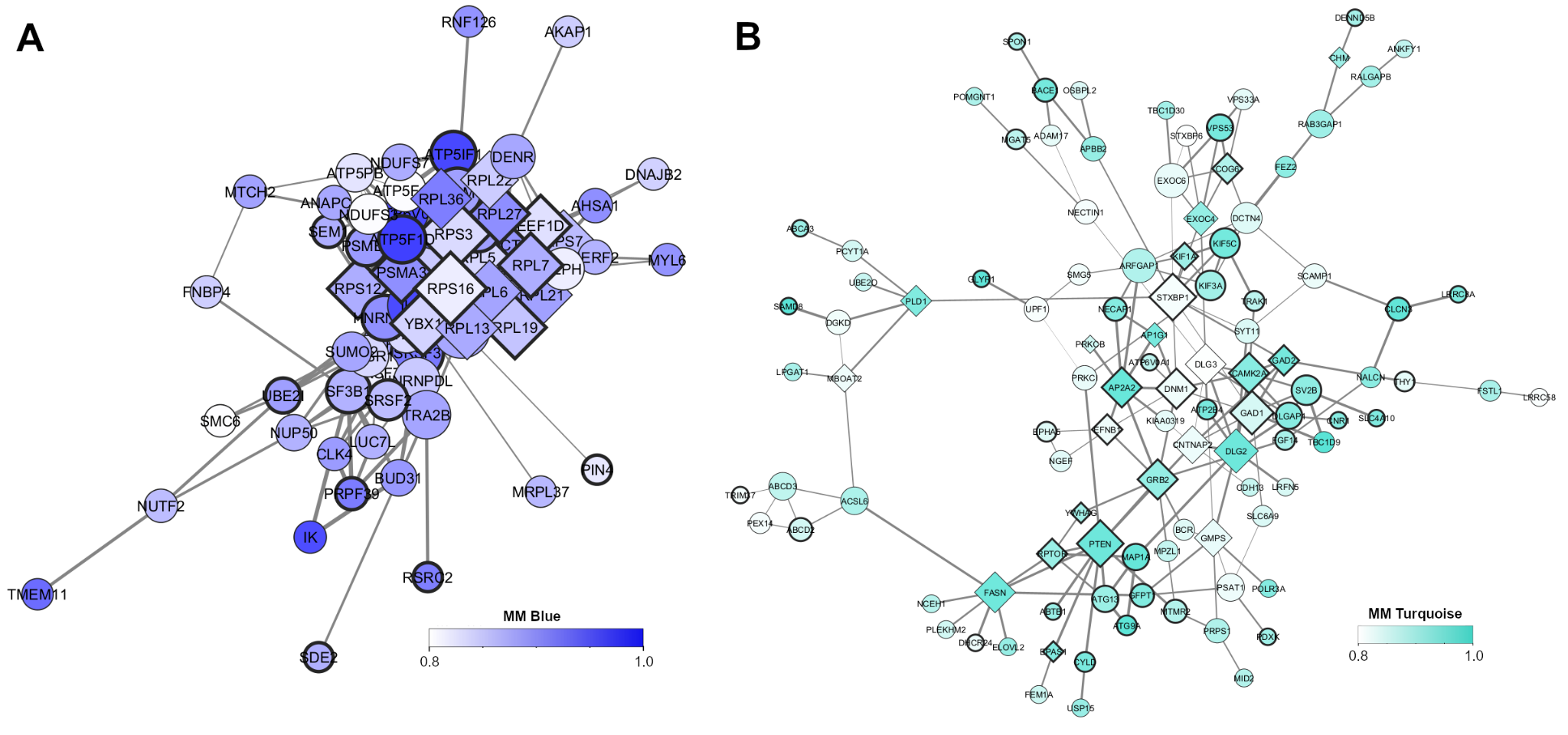

**Figure S7.** Network visualization of the (A) blue and (B) turquoise modules in the caudal hippocampus. Nodes represent genes with significant (p < 0.05) module membership (MM) and gene significance for at least one module-related trait. Node size represents degree and color indicates MM. Nodes with bold borders are differentially expressed between consistently neophobic and non-neophobic sparrows, while diamond-shaped nodes denote PPI hub genes. Edge width reflects WGCNA weights and have been filtered to include only those supported by PPI data.

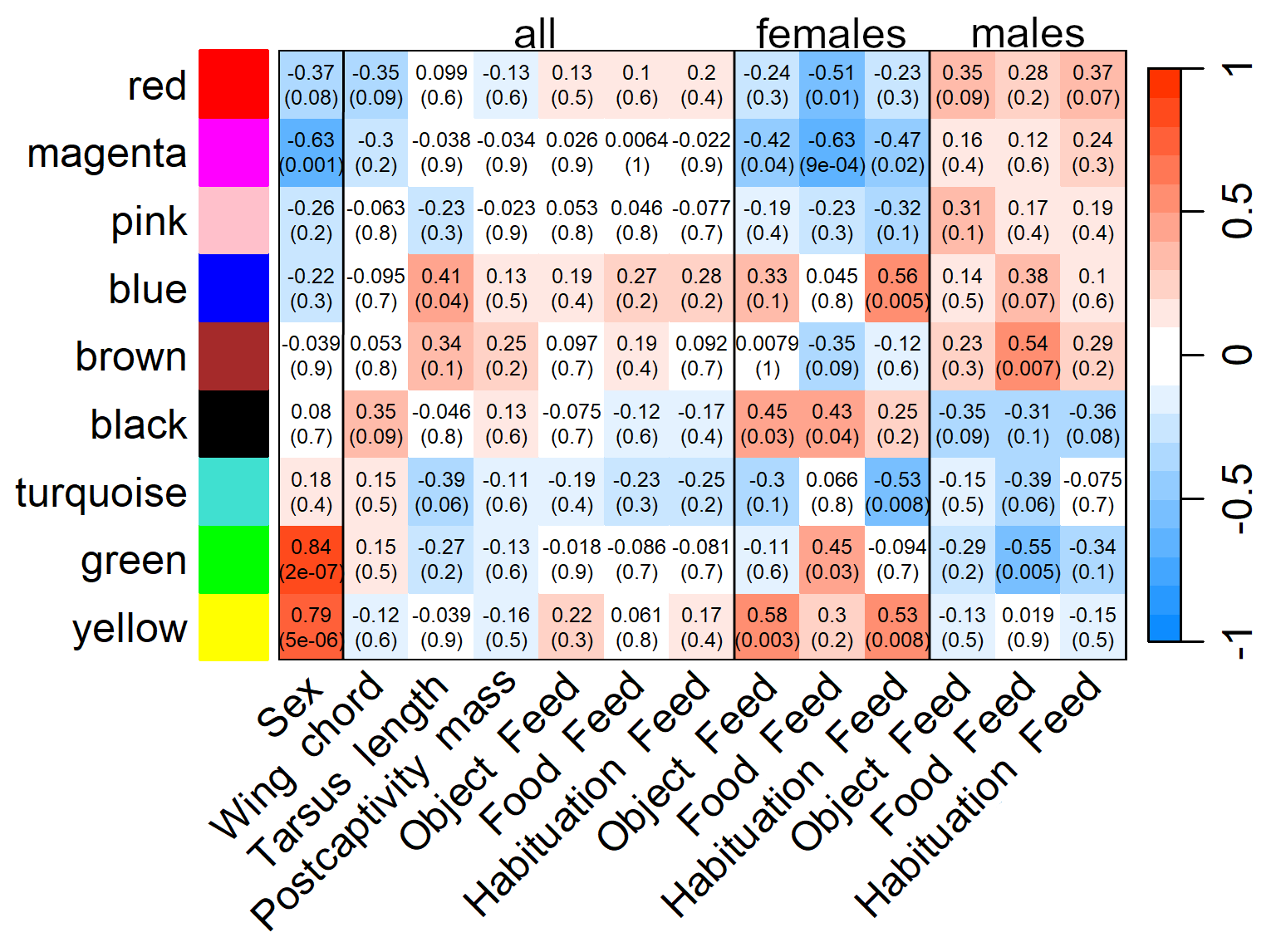

**Figure S8**. Extended module-trait relationships in the sparrow caudal hippocampus were identified using bicor tests in WGCNA. Associations are shown for sex (females as reference), body size (wing chord length, tarsus length, and mass post-captivity), and latency to feed during behavioral trials across and within sexes. “Object Feed” refers to latency to feed from a food dish with a novel object; “Food Feed” from a food dish with a novel food; and “Habituation Feed” from a food dish with the same initially novel object over four days. Correlation coefficients are shown with p-values in parentheses.

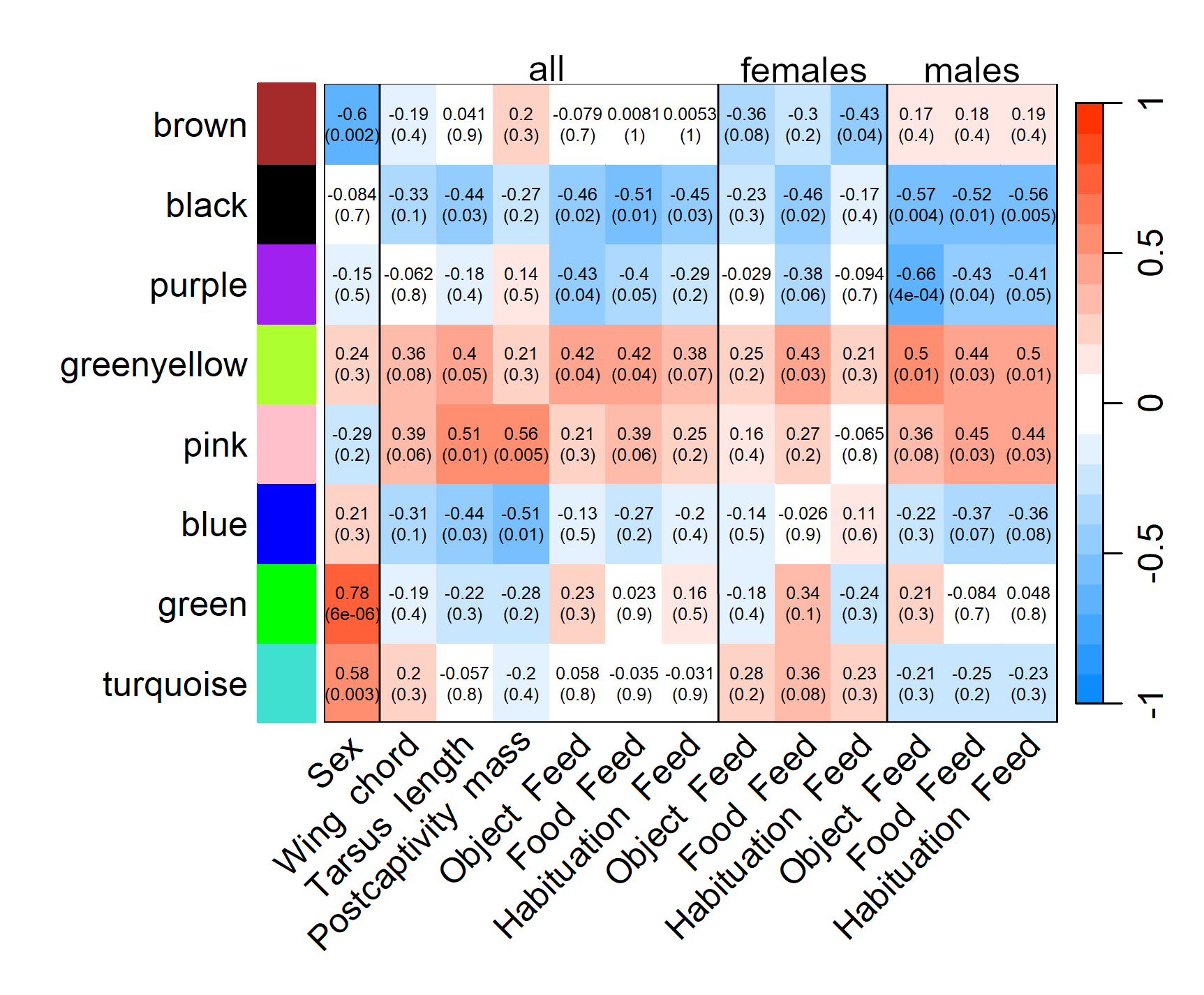

**Figure S9**. Extended module-trait relationships in the sparrow rostral hippocampus were identified using bicor tests in WGCNA. Associations are shown for sex (females as reference), body size (wing chord length, tarsus length, and mass post-captivity), and latency to feed during behavioral trials across and within sexes. “Object Feed” refers to latency to feed from a food dish with a novel object; “Food Feed” from a food dish with a novel food; and “Habituation Feed” from a food dish with the same initially novel object over four days. Correlation coefficients are shown with p-values in parentheses.

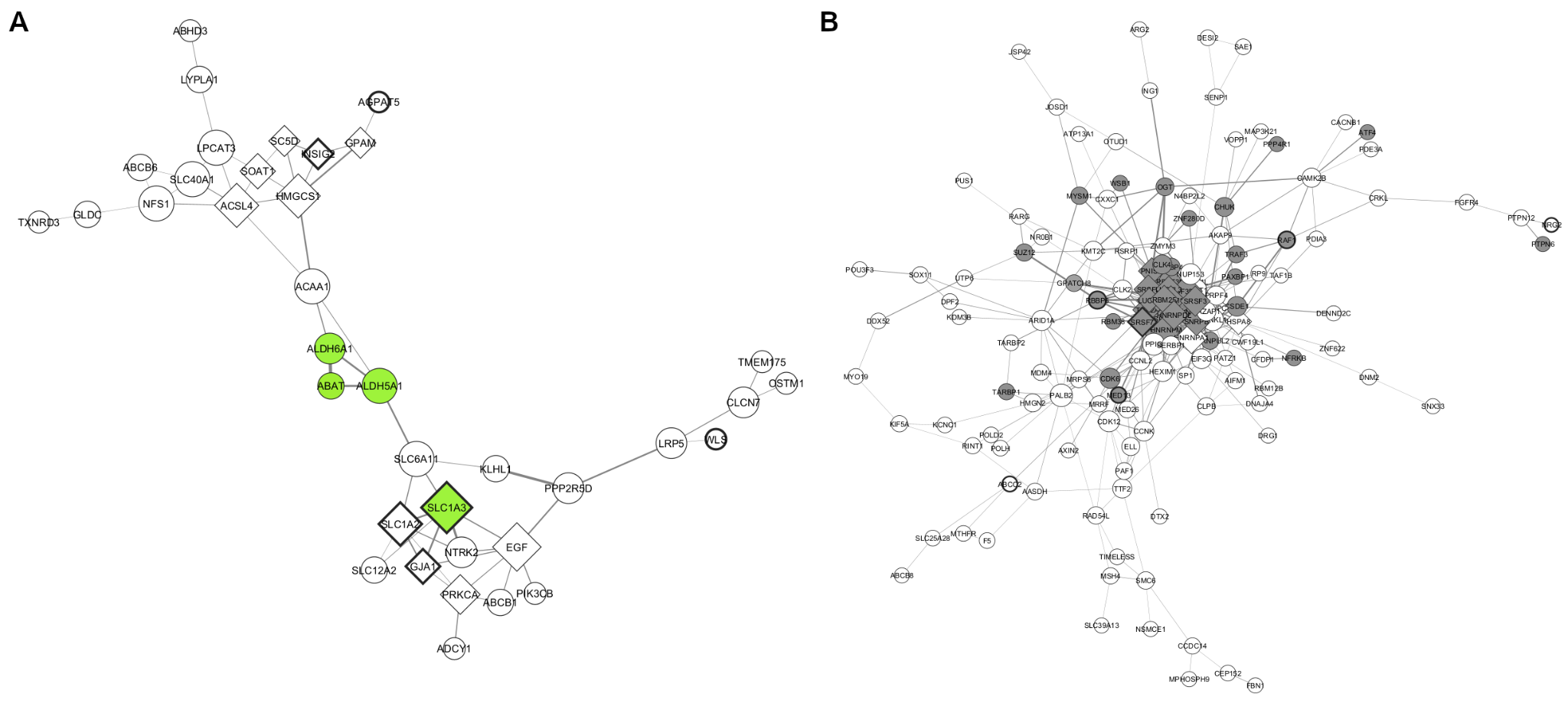

**Figure S10.** Network visualization of the (A) greenyellow and (B) black modules in the rostral hippocampus. Nodes represent genes with significant (p < 0.05) module membership (MM) and gene significance for at least one module-related trait. Node size represents degree and color indicates MM > 0.8. Nodes with bold borders are differentially expressed between consistently neophobic and non-neophobic sparrows, while diamond-shaped nodes denote PPI hub genes. Edge width reflects WGCNA weights and have been filtered to include only those supported by PPI data.

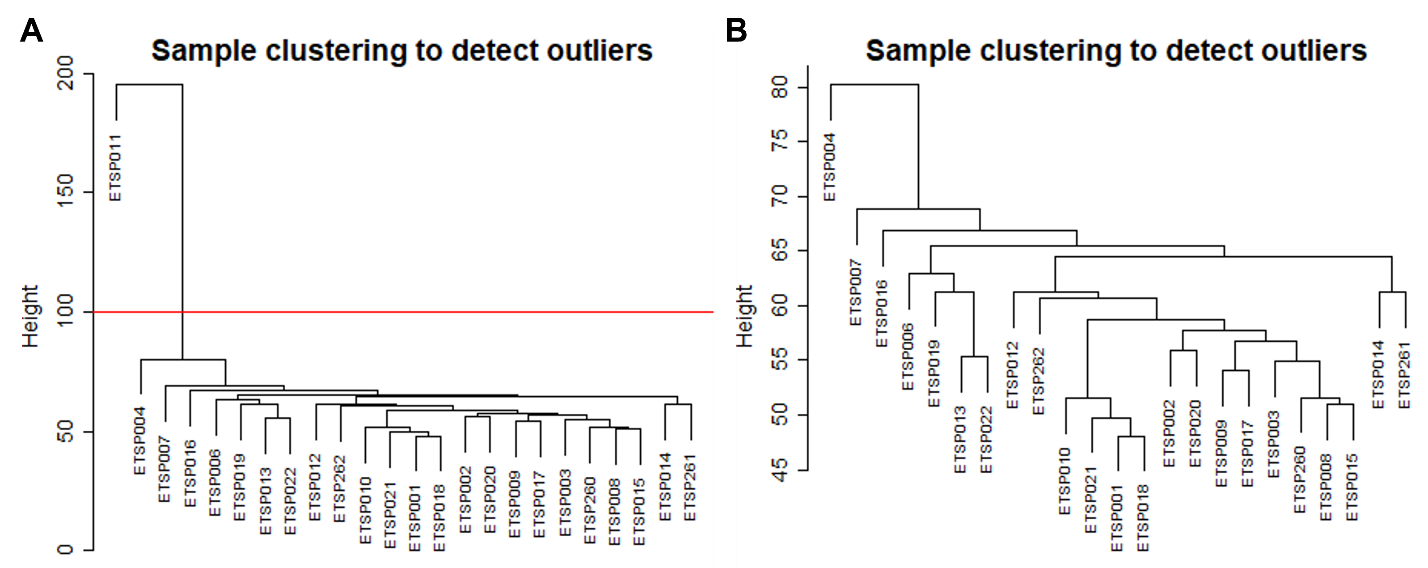

**Figure S11.** Sample clustering to detect outliers in a WGCNA analysis in the striatum (A) before and (B) after outlier removal.

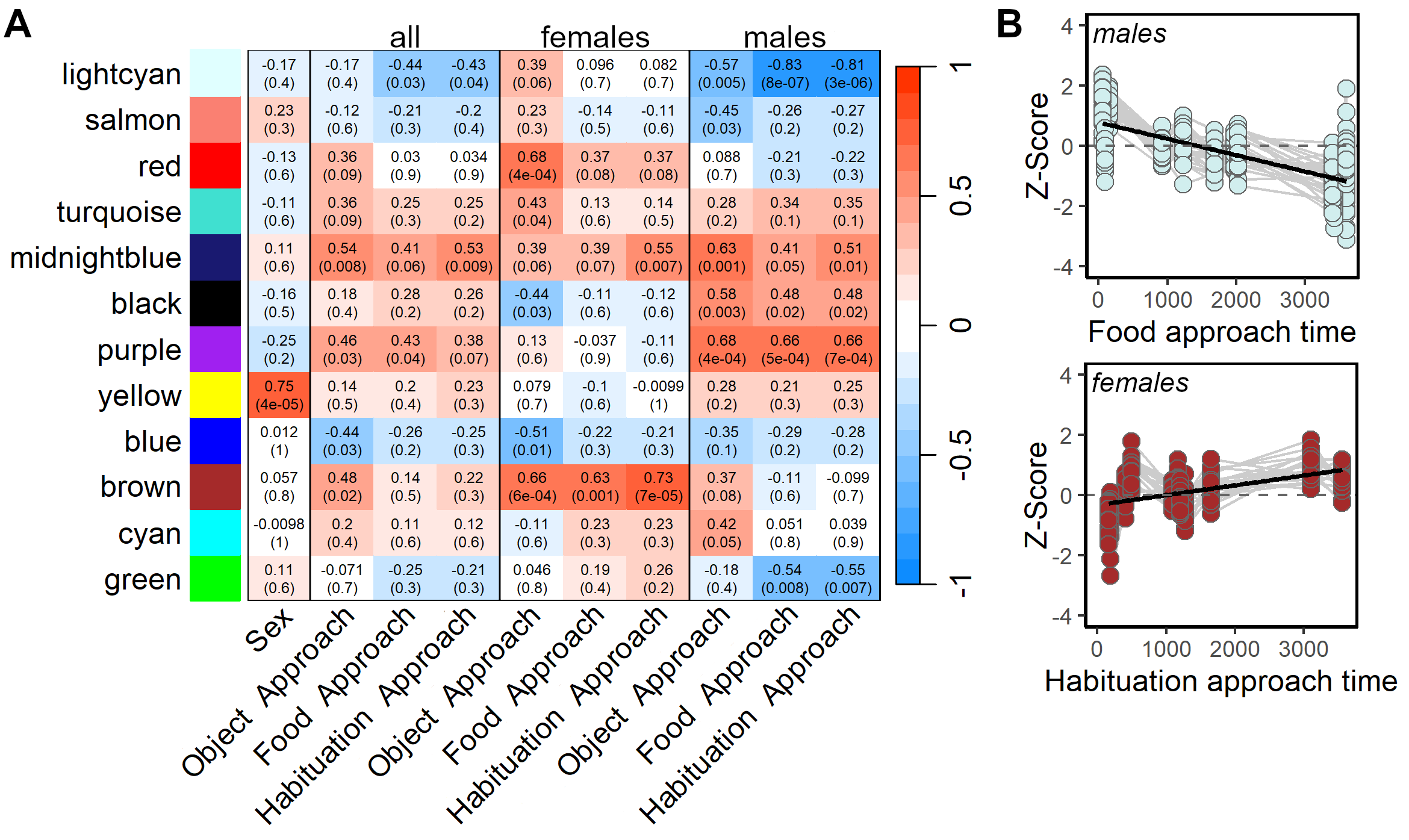

**Figure S12.** (A) Module-trait relationships in the sparrow striatum were identified using bicor tests in WGCNA. Associations are shown for sex (females as reference) and latency to approach in behavioral trials across and within sexes. “Object Approach” refers to the latency to approach a food dish with a novel object; “Food Approach” to a food dish with a novel food; and “Habituation Approach” to repeated exposure to a food dish with the same initially novel object over four days. Correlation coefficients are shown with p-values in parentheses. (B) Relative expression of genes with MM > 0.8 in the light cyan (top) and brown (bottom) modules is shown across food and habituation approach times (in seconds) for males and females. Gray lines connect genes across samples. Z-scores reflect standardized gene expression: Z = (x − μ)/σ, where μ and σ are the mean and standard deviation across all samples and x is the expression of the gene in a specific sample.

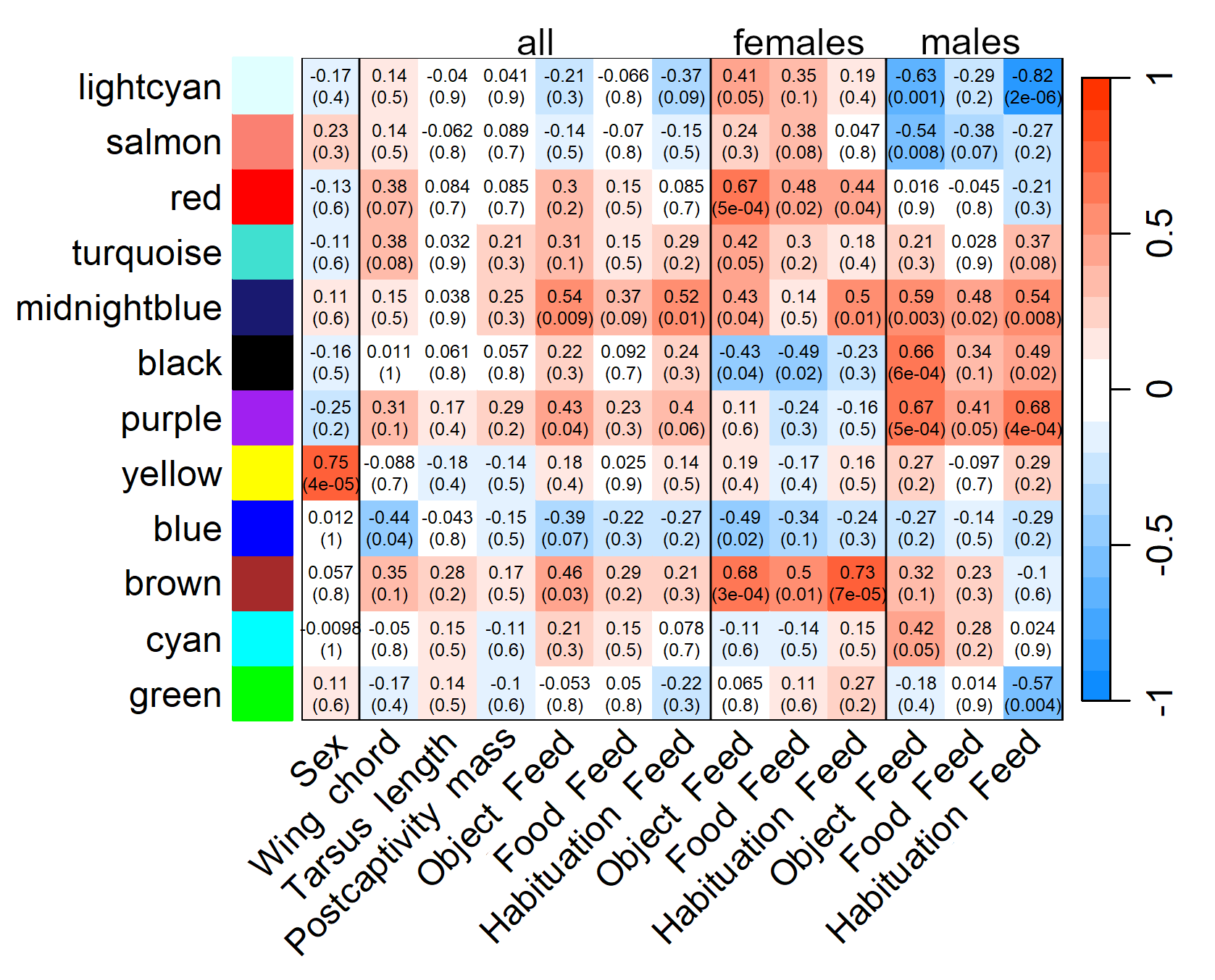

**Figure S13**. Extended module-trait relationships in the sparrow striatum were identified using bicor tests in WGCNA. Associations are shown for sex (females as reference), body size (wing chord length, tarsus length, and mass post-captivity), and latency to feed during behavioral trials across and within sexes. “Object Feed” refers to latency to feed from a food dish with a novel object; “Food Feed” from a food dish with a novel food; and “Habituation Feed” from a food dish with the same initially novel object over four days. Correlation coefficients are shown with p-values in parentheses.

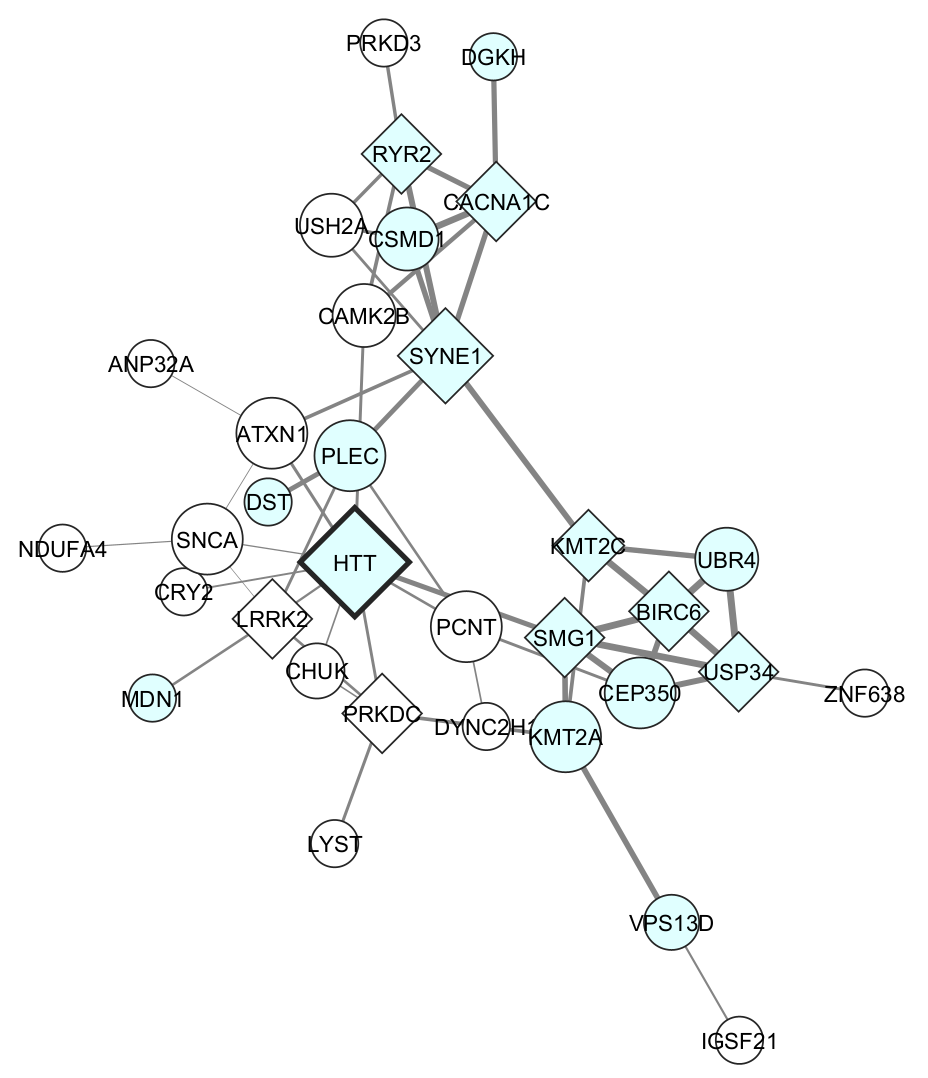

**Figure S14**. Network visualization of the lightcyan module in the striatum. Nodes represent genes with significant (p < 0.05) module membership (MM) and gene significance for at least one module-related trait. Node size represents degree and color indicates MM > 0.8. Nodes with bold borders are differentially expressed between consistently neophobic and non-neophobic sparrows, while diamond-shaped nodes denote PPI hub genes. Edge width reflects WGCNA weights and have been filtered to include only those supported by PPI data.

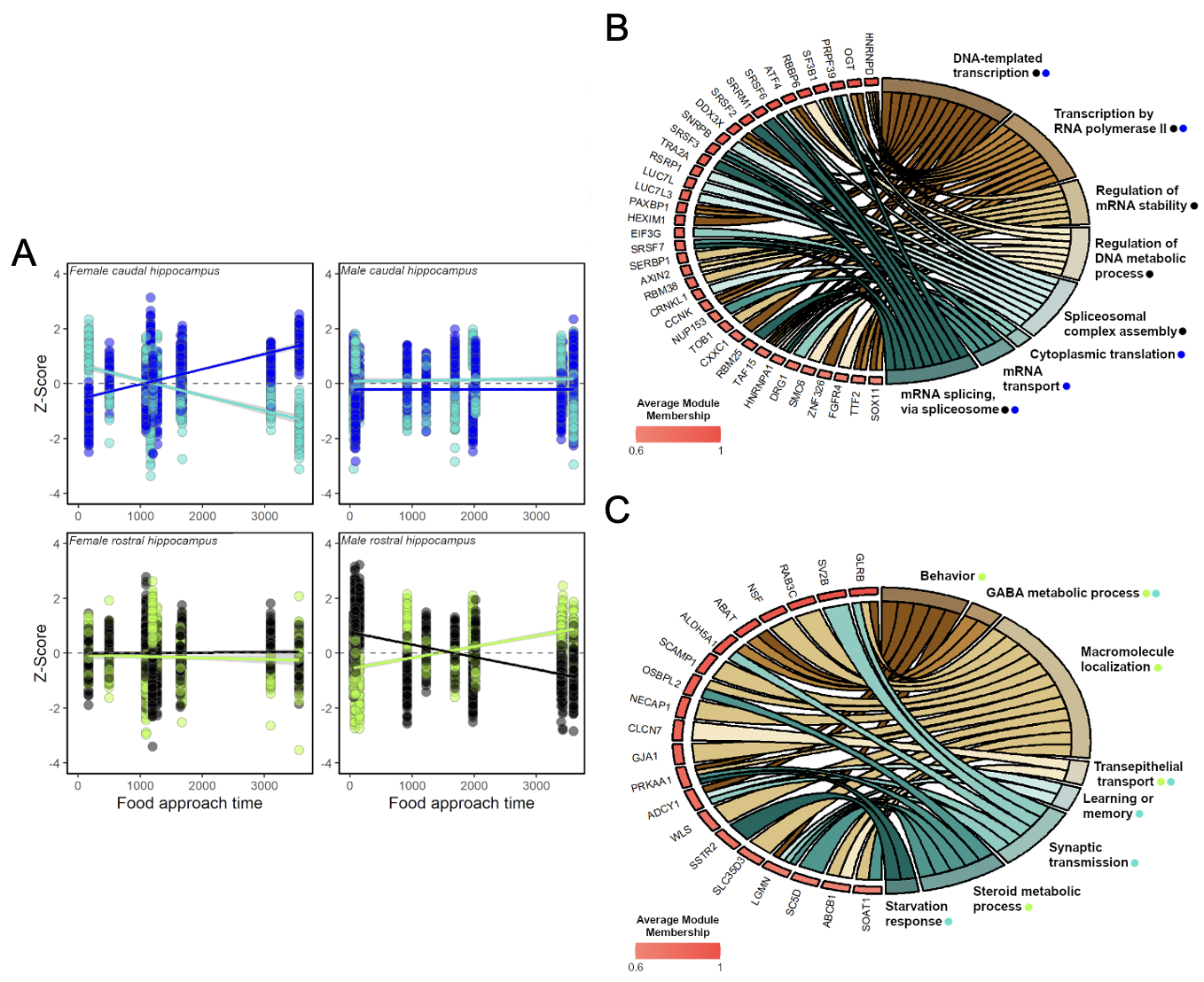

**Figure S15.** Consensus between caudal and rostral hippocampus gene modules in Eurasian tree sparrows. (A) Relationship between relative gene expression (Z-score) and time to approach novel food items (in s) of genes showing overlap in trait-associated modules in females (left panels) and males (right panels). These genes were assigned to the blue/turquoise modules in the caudal hippocampus (top panels) and the greenyellow/black modules in the rostral hippocampus (bottom panels). GOChord plot of overlapping genes in the (B) caudal hippocampus blue module and rostral hippocampus black module and (C) caudal hippocampus turquoise module and rostral hippocampus greenyellow module. All genes have module membership, MM, >0.6 and p<0.05 for novel food approach gene significance in both overlapping modules. Genes are linked to their assigned gene ontology (GO) biological processes via colored ribbons and ordered according to their average MM in overlapping modules. The colored circles next to the GO terms indicate the module the term originated from. Due to similarity, some terms were merged, including: “synaptic transmission” includes both “chemical synaptic transmission” and “positive regulation of synaptic transmission”; “transcription by RNA polymerase II” also includes “positive regulation of transcription by RNA polymerase II”; “DNA-templated transcription” includes “positive regulation of DNA-templated transcription” and “regulation of DNA-templated transcription elongation”; “mRNA splicing, via spliceosome” includes “regulation of mRNA splicing, via spliceosome” and “mRNA cis splicing, via spliceosome”.

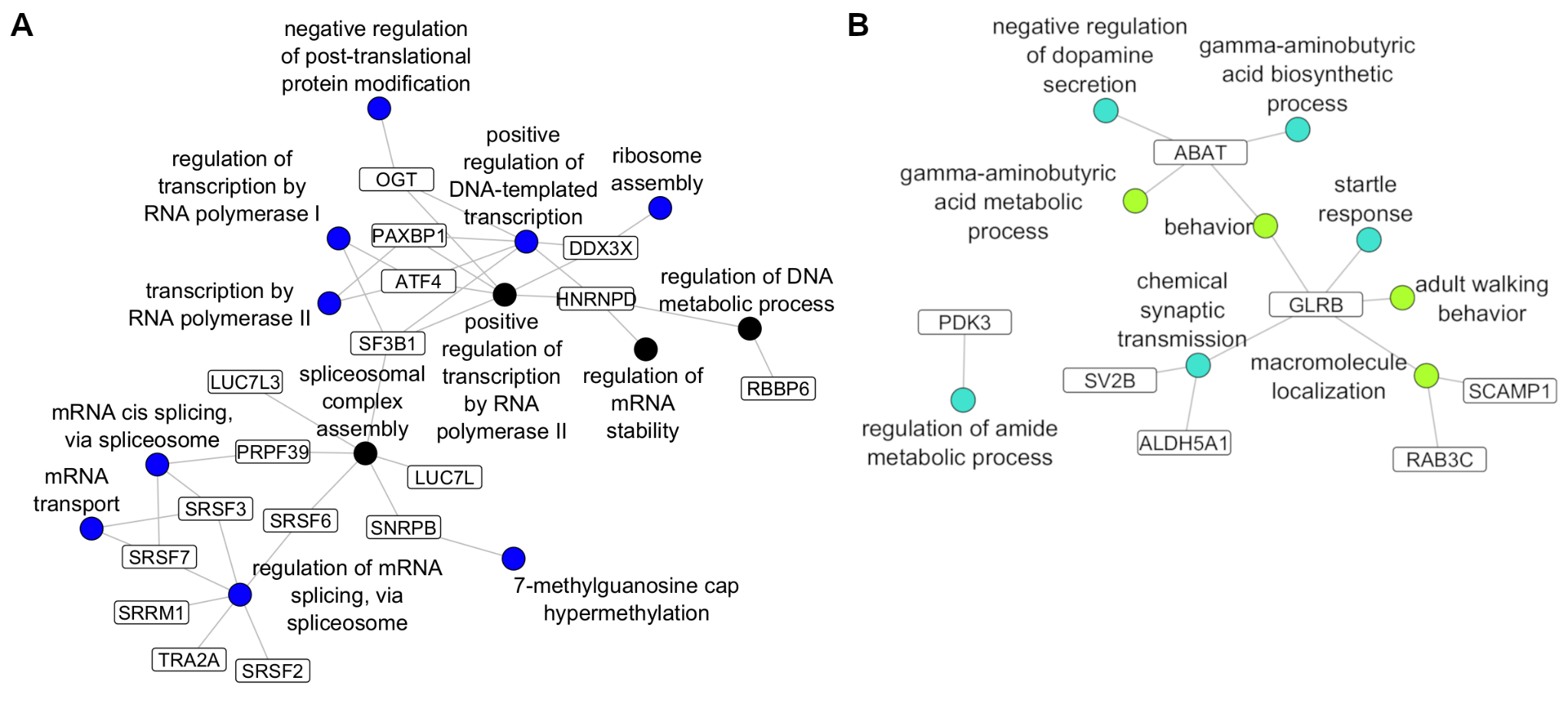

**Figure S16**. Overlap in key genes and Gene Ontology (GO) biological processes associated with Figure 6 (see main text). Shown are genes with high module membership (> 0.80), significant associations with at least one module-related behavioral trait (p < 0.05), and shared presence in corresponding caudal and rostral hippocampus module pairs: (A) blue and black, and (B) turquoise and greenyellow. Gene symbols appear in white boxes and are connected by edges to GO biological process nodes, which are colored according to their module of origin.

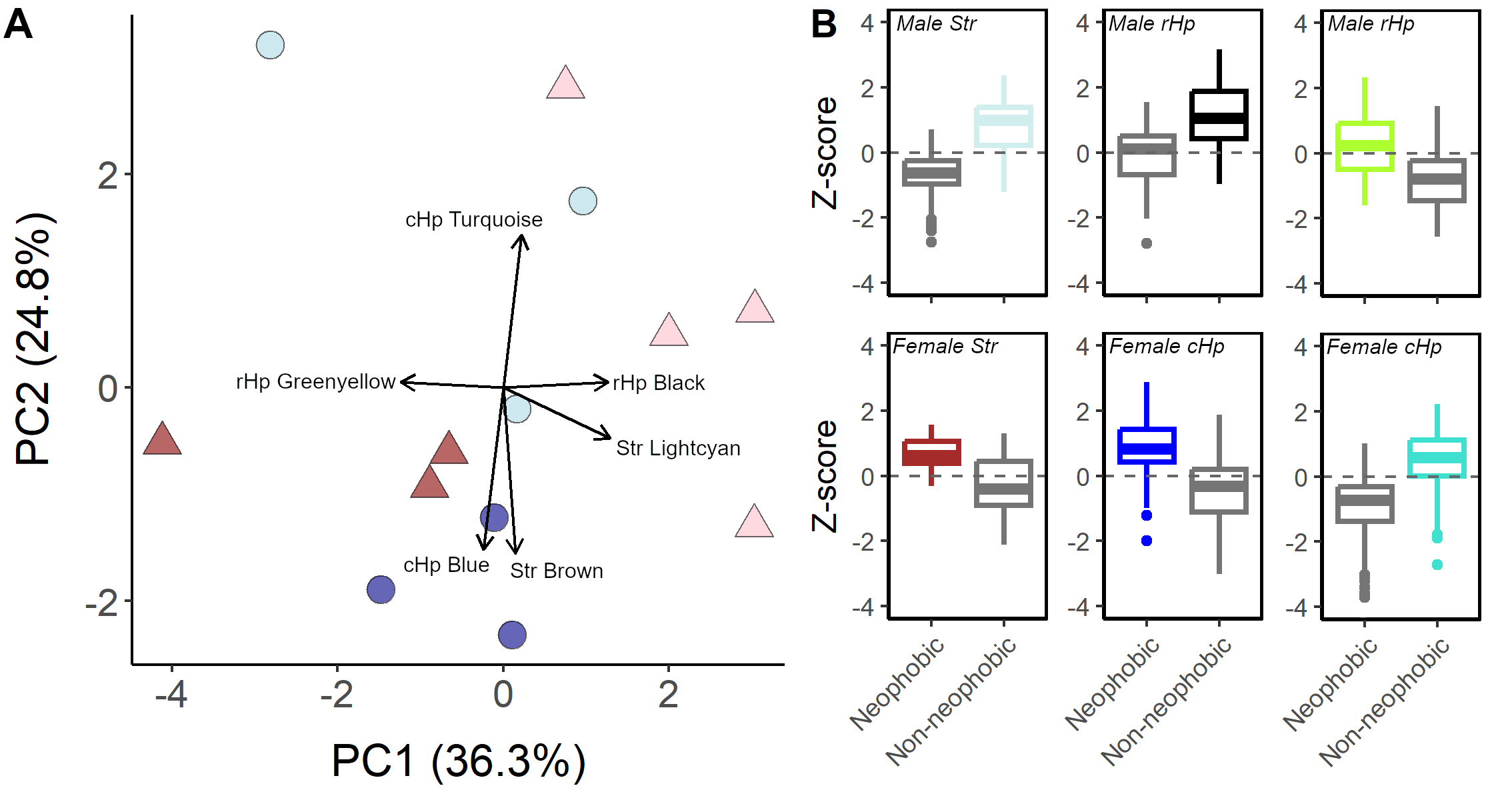

**Figure S17.** (A) Principal component analysis (PCA) visualization of the first two principal components (PC; 61.1% of the variance) of module eigengenes for behavioral trait-related modules from the caudal hippocampus (cHp), rostral hippocampus (rHp), and striatum (Str). Only vectors for PC loadings > |0.4| are shown. Points depict males (triangles, red) and females (circles, blue) that were consistently neophobic (darker shades) or non-neophobic (lighter shades). PC1 loadings: rHp black, 0.423; rHp purple, 0.279; rHp greenyellow, -0.412; cHp magenta, -0.127; cHp blue, -0.079; cHp turquoise, 0.074; Str lightcyan, 0.430; Str red, 0.183; Str midnightblue, -0.231; Str black, -0.398; Str purple, -0.336; Str greenyellow, 0.049. PC2 loadings: rHp black, 0.017; rHp purple, -0.050; rHp greenyellow, 0.017; cHp magenta, 0.005; cHp blue, -0.506; cHp turquoise, 0.476; Str lightcyan, -0.158; Str red, -0.350; Str midnightblue, -0.308; Str black, -0.028; Str purple, -0.045; Str greenyellow, -0.518. (B) Boxplots showing relative gene expression levels (Z-score) for genes from modules with the highest loadings on PC1 (males, top panels) and PC2 (females, bottom panels), across consistently neophobic and non-neophobic sparrows. Box colors indicate module identity and are assigned based on the group (neophobic or non-neophobic) with the highest relative expression.
